# segSHAPE: RNA secondary structure prediction from nanopore direct RNA sequencing

**DOI:** 10.64898/2026.06.15.732177

**Authors:** Guangzhao Cheng, Lassi Härtsiä, Minna-Liisa Änkö, Lu Cheng

## Abstract

RNAs adopt complex structures that regulate key biological processes, making accurate structure prediction essential. Chemical probing coupled with Nanopore direct RNA sequencing (DRS) offers a route to single-molecule structural inference, but current tools are limited by inaccurate signal-to-sequence alignment, which degrades modification-rate estimation and downstream structure prediction. Here we introduce segSHAPE for RNA secondary structure prediction from Nanopore DRS data (both RNA002 and RNA004 chemistries), a probe-agnostic framework that improves signal alignment using prior information of basecalling and per-read signal baseline shift correction, learns position-specific k-mer raw signal parameters, and estimates per-nucleotide modification rates with an unsupervised anomaly detector. On three public RNA002 DRS datasets spanning different chemical probes (AcIm, NAI-N3) and RNAs from 421 to 1552 nt, segSHAPE achieves the highest F1 score and Matthews correlation coefficient (MCC) on all RNAs, exceeding the strongest baseline by 3.4 to 5.8 percentage points in MCC. It additionally captures the ligand-induced conformational change of the thiamine pyrophosphate (TPP) riboswitch RNA directly from RNA002 DRS data using the DEPC probe. On a public RNA004 DRS dataset, segSHAPE improves over the sm-PORE-cupine baseline by 17 ROC-AUC points in modification rate estimation and by 6.7 MCC points in structure prediction. These results establish segSHAPE as a unified, probe-agnostic pipeline for RNA structure prediction from Nanopore DRS data.

## 1 Introduction

RNAs adopt intricate secondary and tertiary structures that regulate diverse stages of their lifecycle, including transcription, splicing, translation, and RNA-protein interactions [1–4]. Accurate RNA secondary structure prediction is therefore essential for elucidating functional mechanisms and has wide implications across molecular biology [5], drug design [6], and synthetic biology [7].

Chemical probing coupled with next-generation sequencing (NGS), exemplified by SHAPE-MaP [8], is currently the most widely used strategy for experimentally inferring RNA structure. A chemical reagent selectively attaches to flexible, single-stranded nucleotides and induces reverse-transcription mutations during NGS library preparation; comparing the mutational profiles of treated and untreated samples yields a per-nucleotide reactivity profile (high for single-stranded regions, low for paired regions) that is then used as a soft constraint to guide secondary structure prediction. Although SHAPE-MaP has substantially advanced the field, it has two intrinsic limitations: PCR amplification artifacts can distort the mutation profile, and short-read sequencing precludes the recovery of full-length, single-molecule structural information [9].

Direct RNA sequencing (DRS) by Oxford Nanopore Technologies (ONT) offers a promising alternative [10]. Probe-induced RNA modifications perturb the nanopore current in a characteristic manner, enabling quantitative detection of modification events without PCR amplification, which makes DRS particularly well suited to structurally complex and long transcripts. Several DRS-based structure-probing pipelines have been developed for RNA002 data, including nanoSHAPE [11], SMS-seq [12], and PORE-cupine [13]. Each combines a chemical probe (AcIm, DEPC and NAI-N3, respectively), a signal-segmentation tool (Tombo [14] or Nanopolish [15]), and a reactivity-scoring strategy (a Kolmogorov-Smirnov test, a hard-constrained modification rate, or a trained support vector machine). More recently, as ONT DRS has moved from RNA002 to the newer RNA004 chemistry, sm-PORE-cupine extended Nanopore structure probing to RNA004 data [16].

Four open problems limit the accuracy and applicability of current DRS-based pipelines. First, the signal-segmentation tools they rely on (Tombo and Nanopolish) use a unified *k*-mer parameter table, implicitly assuming that all positions sharing the same *k*-mer follow identical signal distributions; in practice this assumption breaks down, producing noisy segmentation and misalignment that propagate into modification calling [17, 18]. Second, none of these pipelines exploit basecalling prior information when aligning raw signals to the reference sequence; without such a prior to constrain the search, the dynamic time warping (DTW) and hidden Markov model (HMM) aligners they use leave the signal-to-reference alignment inaccurate near the read ends. Third, a separate source of error is per-read signal baseline drift, which leads to suboptimal signal-to-reference alignment. Fourth, each existing pipeline is tied to a specific chemical probe (e.g. SMS-seq to DEPC), and systematic benchmarking across diverse RNAs and probes is still lacking. A general approach to structure prediction that is probe-agnostic and forward-compatible is becoming increasingly important as novel probes continue to emerge [19, 20].

Here we introduce segSHAPE, a unified computational framework for RNA secondary structure prediction from Nanopore DRS data (both RNA002 and RNA004 chemistries). segSHAPE integrates three complementary design choices that directly address the limitations above: (i) an anchored alignment guided by basecalling prior information from Dorado’s `move` table and complemented by a per-read signal baseline shift correction; (ii) iterative position-specific *k*-mer modeling that estimates a unique signal distribution at each reference position, building on the recently proposed SegPore segmentation [17]; and (iii) per-position modification-rate estimation by an unsupervised anomaly detector. The estimated modification rates are transformed into reactivity scores and passed to RNAfold [21] for centroid structure prediction.

We evaluate segSHAPE on five public DRS benchmark datasets spanning RNA002 and RNA004 chemistries, three chemical probes (AcIm, NAI-N3, DEPC), and RNAs from 165 to 1552 nt (Supplementary Table S1). On RNA002 data, segSHAPE achieves the best F1 and MCC scores across all three benchmark RNAs compared with sequence-only and reactivity-aware baseline methods, and captures the ligand-induced conformational change of the thiamine pyrophosphate (TPP) riboswitch directly from DRS data. On RNA004 data, segSHAPE outperforms sm-PORE-cupine in ROC-AUC for modification-rate estimation and improves downstream structure prediction accuracy.

segSHAPE thus provides a unified, probe-agnostic and forward-compatible pipeline for Nanopore DRS-based RNA secondary structure prediction.

## 2 Results

### 2.1 The segSHAPE pipeline

The segSHAPE workflow is illustrated in Figure 1. Given a control sample, a chemical probe treated sample, and a reference sequence, segSHAPE predicts the RNA secondary structure. The control sample is first used to estimate a position-specific *k*-mer parameter table, which is then fixed and applied to the treated sample to compute per-position modification rates. These modification rates are transformed into reactivity scores and supplied to RNAfold, which converts them into SHAPE-derived pseudo-energies and returns the centroid structure as the final prediction.

**Fig. 1.**
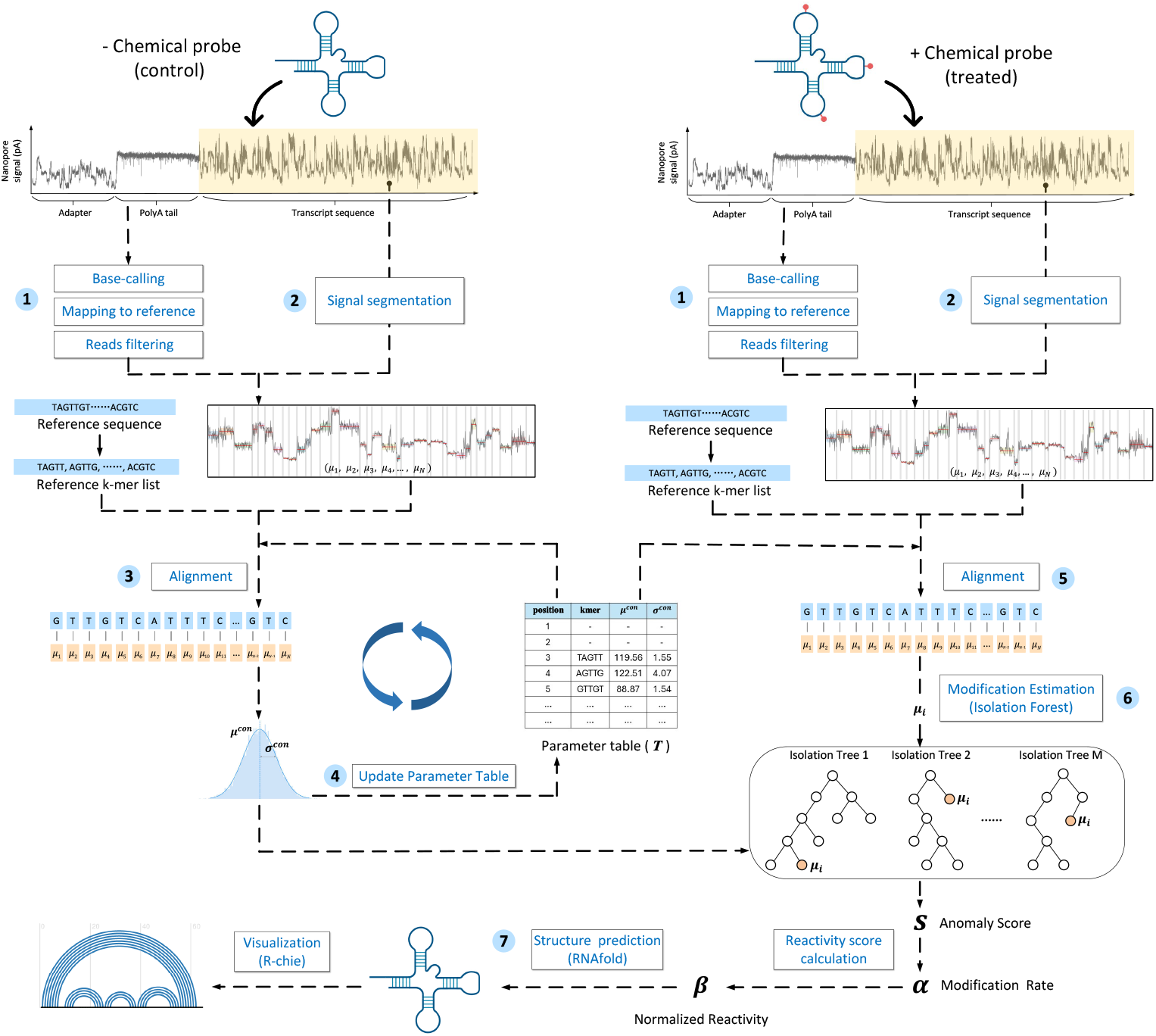
segSHAPE workflow, which requires both a control sample (untreated) and a treated sample (treated with chemical probes). The workflow consists of the following steps: (1) Data preprocessing for both samples; (2) Signal segmentation for both samples, resulting in signal segments (sub-events); (3) Alignment of control sample sub-events to the reference sequence, guided by basecalling-derived anchor points and complemented by a per-read baseline shift correction, returning an event (merged sub-events) per reference position per read; (4) Estimation of the position-specific *k*-mer parameter table from the control sample events, with iterative refinement by repeating steps 3-4; (5) Alignment of treated sub-events using the estimated position-specific *k*-mer parameter table; (6) Per-position modification rate estimation by an unsupervised anomaly detector, with Isolation Forest as the default and Gaussian mixture model as an interchangeable alternative, resulting in a modification rate per position per read; (7) RNA secondary structure prediction with RNAfold, with reactivity scores transformed from the modification rates.

The accuracy of any DRS-based structure prediction pipeline depends critically on upstream signal alignment and modification calling. segSHAPE improves these steps through three complementary design choices: an anchored alignment guided by basecalling prior information; a position-specific *k*-mer model that replaces the unified *k*-mer assumption used by Nanopolish and Tombo; and unsupervised per-position modification-rate estimation with interchangeable outlier detectors. Each component is generally described below and detailed in the Methods section.

#### 2.1.1 Anchored alignment with basecalling prior

Recent advances in the basecaller Dorado provide a `move` table that yields per-read anchor points in both the raw signal and the reference sequence. Anchoring signal alignment on such a basecaller `move` table, rather than running an unconstrained search, has been shown to be faster and more robust than adaptive-banding aligners [22]; segSHAPE adopts this prior and tailors it to SHAPE structure probing. Existing DRS-based structure-probing pipelines (nanoSHAPE, SMS-seq, PORE-cupine) instead align raw signals to the reference without a basecalling prior, which leaves them disadvantaged in three respects: (i) Without the basecalling prior, the transcript region signal might be less accurate due to unreliable poly(A)/adapter localization (see the poor poly(A) detection example by Nanopolish in Supplementary Figure S1). Either part of the poly(A) tail signal might leak into the transcript-region signal, or part of the 3^*′*^ end signal of the transcript might get truncated. (ii) Due to the potentially mis-specified transcript-region boundaries, the dynamic time warping (DTW) and hidden Markov model (HMM) aligners they use generate less accurate signal-to-reference alignment in general. (iii) There exists per-read signal baseline drift, which are not corrected in the existing pipelines and leads to suboptimal signal-to-reference alignment.

segSHAPE addresses these disadvantages as follows. (i) Using Dorado, we derive anchor points in both the raw signal and the reference sequence from the basecaller’s `move` table; these are used as soft constraints in a modified global dynamic-programming alignment algorithm that compares admissible start/end combinations within per-read entry and exit boxes (Supplementary Figure S2) and returns the best-scoring combination. The transcript-region signal is delimited directly from Dorado’s `ts` and `ns` BAM tags together with the `move` table. (ii) To improve the alignment further, segSHAPE seeks a per-read baseline shift *β* that minimizes the average deviation between aligned event means and the corresponding position-specific reference means, over all positions in the transcript. This is done iteratively until the alignment log-likelihood changes by less than 0.1 between successive rounds.

#### 2.1.2 Position-specific *k*-mer modeling

Current DRS signal-alignment tools such as Nanopolish and Tombo rely on a unified *k*-mer parameter table, implicitly assuming that all positions sharing the same *k*-mer follow identical signal distributions. This assumption is inconsistent with what we observed in real data.

As shown in Figure 2A, five distinct positions in pri-miR-17∼ 92 (positions 247, 405, 502, 708 and 911) all carry the same *k*-mer “TTTTG” yet exhibit visibly different empirical signal distributions estimated from Nanopolish’s events, even though Nanopolish models them with a single shared distribution given by the ONT *k*-mer reference (dashed line); the same is observed for a second *k*-mer (“AAGGT”) at five positions in *B. subtilis* 16S rRNA (Figure 2B). A direct consequence of this mismatched assumption is signal misalignment: at position 91 of pri-miR-17 ∼ 92 (*k*-mer “AAGAT”), for instance, Nanopolish yields a bimodal distribution in the control sample (Figure 2C, top), whose two peaks closely match the distributions of its neighboring positions 90 and 92. Because every read in the control sample is by construction unmodified, each position should follow an unimodal distribution (i.e. a single peak); the observed bimodality therefore reflects alignment errors in which signals from positions 90 or 92 might be pulled into position 91.

**Fig. 2.**
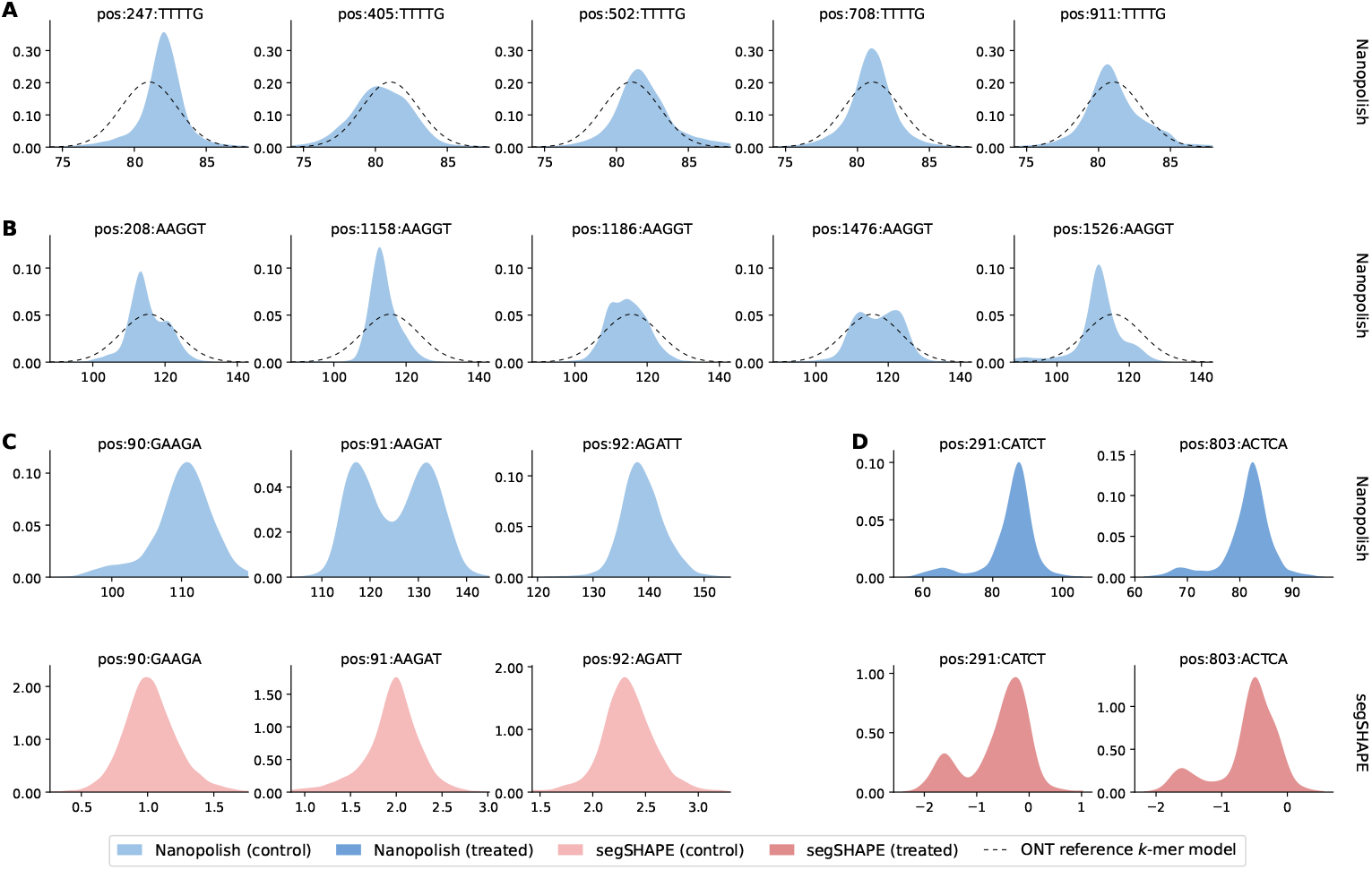
Per-position *k*-mer current signal distributions from Nanopolish and segSHAPE. Filled curves are empirical kernel density estimates (Nanopolish in blue, segSHAPE in pink); dashed line represents the ONT reference *k*-mer model. (A, B) The same *k*-mer at five distinct positions of the pri-miR-17 ∼ 92 (A: TTTTG at positions 247, 405, 502, 708, 911) and *B. subtilis* 16S rRNA (B: AAGGT at positions 208, 1158, 1186, 1476, 1526) control samples shows visibly different signal distributions based on Nanopolish’s alignment. The ONT model is identical across the five sub-panels in each row by construction and serves as a fixed reference. (C) Three consecutive positions (positions 90, 91, 92; GAAGA / AAGAT / AGATT) in the pri-miR-17 ∼ 92 control sample. Top (Nanopolish): position 91 shows a bimodal distribution; bottom (segSHAPE): position-specific alignment recovers the expected unimodal distribution. (D) Two chemically-modified positions (position 291 CATCT, position 803 ACTCA) in the pri-miR-17∼ 92 treated sample. Compared with Nanopolish (top), segSHAPE (bottom) provides clearer separation between the unmodified signal and the modified component, which appears as a small left-shifted peak.

segSHAPE replaces the unified *k*-mer table with a position-specific parameter table that estimates an independent set of parameters (mean and variance) for each reference position, even when multiple positions share the same *k*-mer. The table is initialized from ONT’s reference *k*-mer model and then iteratively refined over three rounds: in each round, control reads are aligned using the table estimated from the previous round, and then the parameters of every position are re-estimated from its aligned events (Figure 1, steps 3 and 4). After refinement, position 91 in segSHAPE recovers the expected unimodal distribution (Figure 2C, bottom) in the control sample.

#### 2.1.3 Modification detection and reactivity scoring

The position-specific parameter table is estimated from the control sample and fixed for the alignment of the treated sample. For each position in the reference sequence, we build a null distribution of signal baselines from the sub-events aligned to the position in the control sample. We then apply an unsupervised anomaly detector to sub-events aligned to the position in the treated sample to identify the outliers using this null distribution, and quantify the modification rate as the proportion of outliers.

Several anomaly detectors can be used for this purpose, including Isolation Forest [23], Gaussian mixture model [24], One-Class SVM [25], the absolute difference of medians (|Δmedian|), the Kolmogorov-Smirnov two-sample statistic [26], and the 1-D Wasserstein distance [27] and xPore [28]. We performed an ablation study of the seven detectors in three RNA002 benchmark datasets and found that Isolation Forest and GMM have the best performances (Supplementary Table S3). As Isolation Forest is distribution-free, it imposes the weakest assumptions on the control signal. We use Isolation Forest as the default anomaly detector, but also include GMM as an interchangeable detector. The derived modification rates are then normalized to a standard normal distribution (z-score) to obtain the reactivity scores, which are used as input to RNAfold to predict the secondary structure.

### 2.2 Case study: pri-miR-17∼92

As an initial validation, we tested segSHAPE on the pri-miR-17 ∼ 92 RNA, a 951-nt primary transcript that encodes six oncogenic microRNAs (miR-17, miR-18a, miR-19a, miR-20a, miR-19b and miR-92a) and folds into a corresponding set of six independent hairpins. The original nanoSHAPE paper [11] provide both Nanopore DRS RNA002 SHAPE data and SHAPE-MaP data as the ground truth for the pri-miR-17 ∼ 92 RNA, with 1-acetylimidazole (AcIm) as the chemical probe. We used the SHAPE-MaP data to benchmark the per-position reactivity and the predicted secondary structure of segSHAPE.

The centroid structure predicted by segSHAPE recovers all six conserved miRNA hairpins, in agreement with the reference SHAPE-MaP structure (Figure 3B). The arc-diagram representation shows that both methods identify the same set of helical stems for each hairpin, with the per-hairpin base-pair patterns and overall arrangement broadly consistent between the two methods; the spans of the six canonical pre-miRNA hairpins (miR-17, miR-18a, miR-19a, miR-20a, miR-19b and miR-92a) are highlighted as red double-headed arrow lines. This demonstrates that segSHAPE reproduces the canonical secondary-structure of pri-miR-17∼ 92 directly from Nanopore DRS data.

**Fig. 3.**
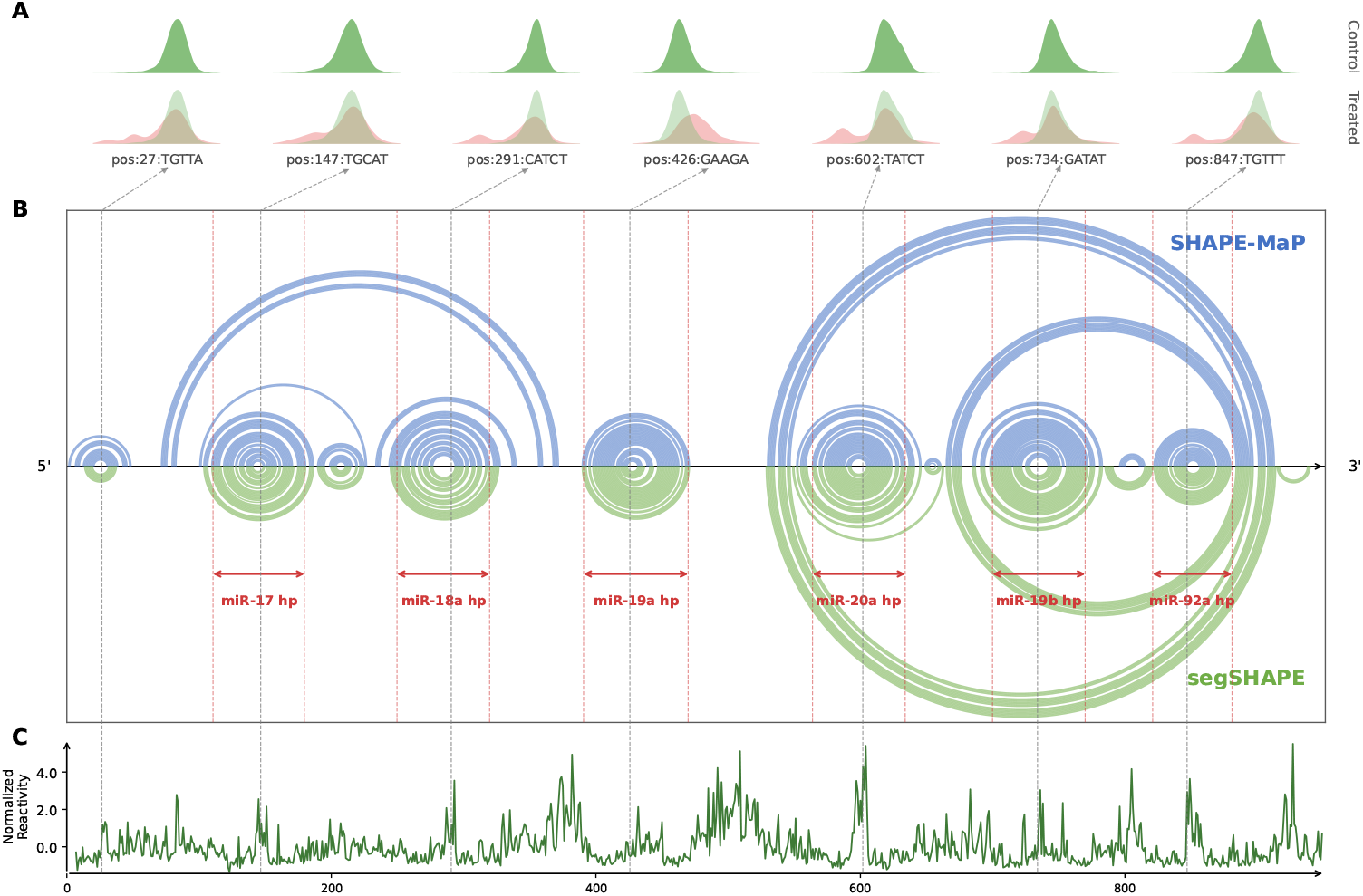
Structural prediction of pri-miR-17 ∼ 92 RNA. Vertical dashed lines connect the seven representative positions across all three panels. (A) segSHAPE signal baseline density plots at seven representative positions, drawn from the loops of a short 5^*′*^-proximal hairpin (position 27) and the six conserved pre-miRNA hairpins miR-17 (147), miR-18a (291), miR-19a (426), miR-20a (602), miR-19b (734) and miR-92a (847). Top row: control sample distribution (no chemical probe, green); bottom row: treated sample distribution (pink) overlaid with the control distribution (light green) so that the modification component is visible as the pink shape that deviates from the control distribution. (B) Arc-diagram comparison of the SHAPE-MaP secondary structure (upper, blue) and the segSHAPE centroid structure prediction (lower, green) along the 951-nt reference. The black central arrow line runs 5^*′*^ → 3^*′*^ on the reference sequence; arc height is rescaled vertically by a constant factor for clarity. Below the central arrow line, red double-headed arrow lines mark six conserved miRNA hairpins (hp): miR-17, miR-18a, miR-19a, miR-20a, miR-19b and miR-92a. (C) segSHAPE normalized SHAPE reactivity profile (darker green line) along the reference sequence; the *x*-axis is shared with Panel B.

At the single-nucleotide level, the modification calls are consistent with the expected SHAPE chemistry. Seven representative positions are drawn from each of the six conserved miRNA hairpin loops plus one short 5^*′*^-proximal hairpin loop (Figure 3A), each show a clear modification component in the treated sample (pink) that is absent from the unmodified control (green), as expected for single-stranded loop bases that are preferentially modified by AcIm. This pattern is observed transcript-wide in the per-position reactivity profile (Figure 3C), with elevated reactivity concentrated in the loop regions and suppressed reactivity in the double-stranded stem regions.

Together, these results show that segSHAPE delivers both a consistent structure with SHAPE-MaP and the expected qualitative SHAPE modification pattern at single-nucleotide resolution.

### 2.3 Structure prediction benchmark on RNA002 datasets

We next benchmarked segSHAPE against six baseline methods on three public Nanopore DRS RNA002 datasets using distinct chemical probes: pri-miR-17∼ 92 (951 nt, AcIm) [11], *Tetrahymena* preribosomal RNA (421 nt, NAI-N3), and *Bacillus subtilis* 16S rRNA (1552 nt, NAI-N3) [13]. The *Tetrahymena* preribosomal intron is the first identified ribozyme and has a well-characterized secondary structure [29], and 16S rRNA has been extensively studied for its highly conserved secondary and tertiary structures, making both ideal benchmarks for RNA structure prediction. The ground truth structure of all three RNAs are provided in the original publications.

Six baseline methods are used for the benchmark: three sequence-only predictors (MXfold2 [30], RNAPKplex [21], RNAfold [21]), and three reactivity-aware DRS pipelines (SMS-seq [12], nanoSHAPE [11], PORE-cupine [13]). segSHAPE is evaluated with either Isolation Forest (IF) or Gaussian mixture model (GMM) as the per-position modification detector. MXfold2 has a maximum sequence length limit of 1000 nt, making it incapable of predicting the *Bacillus subtilis* 16S rRNA (1552 nt).

The ground truth structure is converted into a list of pairs for paired positions (positive cases), and a list of pairs for unpaired positions (negative cases). The same is done for the predicted structures. Then a confusion matrix is constructed, and the following metrics are calculated from the confusion matrix: precision, recall, F1 score, and Matthews correlation coefficient (MCC). See Supplementary Figure S5 for the detailed calculation.

As shown in Table 1, both segSHAPE variants achieve the best F1 and MCC among all methods on every benchmark RNA. We use MCC as the primary basis for comparison because, among the four metrics, it is the only one that jointly accounts for all four confusion-matrix categories and remains reliable under the paired/unpaired class imbalance intrinsic to base-pair classification [31]. By MCC, both segSHAPE variants outperform every baseline on all three RNAs, and the stronger variant exceeds the strongest baseline by 3.4, 4.0 and 5.8 MCC points on pri-miR-17 ∼ 92, *Tetrahymena* and 16S rRNA, respectively. Because the two variants share the entire upstream pipeline (anchored alignment with basecalling prior, position-specific *k*-mer modeling and per-read baseline shift correction) and differ only in the final per-position outlier detector, their consistent gains suggest that improved upstream signal alignment and estimation of the unmodified per-position signal distribution are major contributors to the performance improvement.

**Table 1.** Benchmark results of structure prediction on RNA002 data. P, precision; R, recall; F1, F1 score; MCC, Matthews correlation coefficient (all in percentage). The two segSHAPE rows differ only in the per-position modification detector (GMM vs. Isolation Forest). MCC, which jointly accounts for all four confusion-matrix categories and is robust to class imbalance, is the primary comparison metric; both segSHAPE variants achieve the highest MCC on every RNA. **Bold** number marks the best performance in each column; underlined number marks the second best.

| Model | Reactivity | pri-miR-17~92 RNA |  |  |  | Tetrahymena RNA |  |  |  | Bacillus subtilis 16S rRNA |  |  |  |
| --- | --- | --- | --- | --- | --- | --- | --- | --- | --- | --- | --- | --- | --- |
|  |  | P | R | F1 | MCC | P | R | F1 | MCC | P | R | F1 | MCC |
| MXfold2 | ✗ | 53.62 | 62.21 | 57.60 | 57.73 | 68.66 | **74.19** | 71.32 | 71.33 | - | - | - | - |
| RNAPKplex | ✗ | 66.56 | 82.06 | 73.50 | 73.89 | 52.59 | 57.26 | 54.83 | 54.81 | 53.94 | **58.30** | 56.03 | 56.06 |
| RNAfold | ✗ | 66.87 | <u>83.21</u> | 74.15 | 74.58 | 61.87 | 69.35 | 65.40 | 65.45 | 49.52 | <u>54.47</u> | 51.87 | 51.91 |
| SMS-seq | ✓ | 85.48 | 78.63 | 81.91 | 81.97 | 70.09 | 60.48 | 64.94 | 65.07 | 48.75 | 37.45 | 42.36 | 42.71 |
| nanoSHAPE | ✓ | 85.11 | 76.34 | 80.48 | 80.59 | 62.65 | 41.94 | 50.24 | 51.20 | 63.01 | 42.77 | 50.95 | 51.89 |
| PORE-cupine | ✓ | 85.32 | 82.06 | 83.66 | 83.66 | 81.00 | 65.32 | 72.32 | 72.71 | 61.79 | 48.51 | 54.35 | 54.73 |
| segSHAPE (GMM) | ✓ | <u>85.71</u> | 82.44 | <u>84.05</u> | <u>84.05</u> | **83.50** | 69.35 | <u>75.77</u> | <u>76.07</u> | **70.44** | 54.26 | **61.30** | **61.81** |
| segSHAPE (IF) | ✓ | **89.84** | **84.35** | **87.01** | **87.04** | <u>83.02</u> | <u>70.97</u> | **76.52** | **76.73** | <u>65.12</u> | 50.85 | <u>57.11</u> | <u>57.53</u> |

A second observation is that reactivity inputs do not automatically improve structure prediction. On *Tetrahymena* and 16S rRNA, two of the three reactivity-aware baseline methods (SMS-seq and nanoSHAPE) perform *worse* than the reactivity-free baseline method RNAfold by F1 and MCC. This suggests that low-quality reactivity profiles can actually degrade prediction when upstream signal alignment is noisy. segSHAPE avoids this regression by improving the reactivity signal at its source, through position-specific *k*-mer modeling and basecalling-prior integration. As a result, it delivers consistent gains across all three RNAs and both chemical probes tested.

### 2.4 Dynamical structure estimation of TPP-riboswitch RNA

Beyond static structure prediction, we asked whether segSHAPE can resolve a *dynamic* structural rearrangement directly from Nanopore DRS data. Riboswitches are well suited to this question because they are cis-acting regulatory RNAs that adopt distinct secondary structures in their ligand-free and ligand-bound states. The thiamine pyrophosphate (TPP, 165 nt) riboswitch is among the best characterized [32] and is widely conserved across bacteria, fungi and plants, where it regulates thiamine biosynthesis and transport genes. Because ligand binding triggers a defined rearrangement of the aptamer domain that ultimately controls downstream translation, recovering this conformational change from chemical-probing DRS data provides a focused test of our signal-processing pipeline.

The SMS-seq study [12] provides RNA002 DRS data for the TPP riboswitch in both ligand-free and TPP-bound states, using Diethyl pyrocarbonate (DEPC) as the chemical probe. We applied segSHAPE to both datasets and qualitatively compared the resulting structures with the corresponding references (Figure 4) and predicted structures of SMS-seq. Due to the low read coverage in this TPP-riboswitch dataset, two parameters in the Isolation Forest outlier detector were re-tuned in segSHAPE to compensate for the low read coverage (details in Supplementary Methods Section 4.1). In the ligand-free state (Figure 4A), segSHAPE reproduces the reference aptamer topology, whereas SMS-seq miscalls several stretches of pairings and yields a noticeably different structure. In the TPP-bound state (Figure 4B), segSHAPE again captures the structure change (highlighted in circle) like a “clamp” holding TPP in the center, which is missed by SMS-seq.

**Fig. 4.**
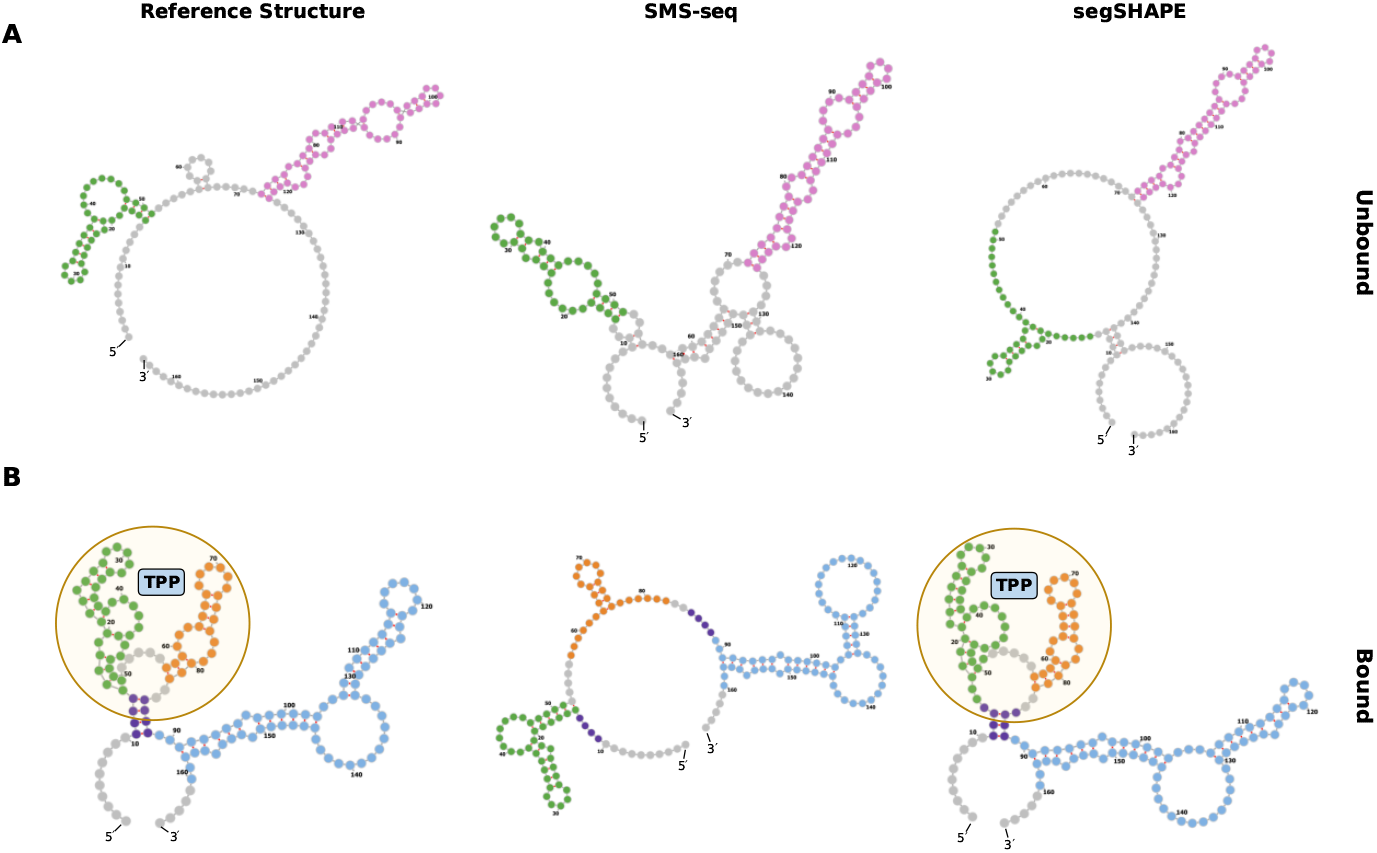
segSHAPE captures TPP-riboswitch structural dynamics. RNA secondary structures of the thiamine pyrophosphate (TPP) riboswitch in (A) the unbound state and (B) the TPP-bound state, comparing the reference secondary structure, predicted structures by SMS-seq and segSHAPE. segSHAPE tracks the ligand-induced conformational change (highlighted by the circle in panel B), whereas SMS-seq does not.

Two observations follow from this comparison. First, ligand-induced riboswitch conformational dynamics can be recovered directly from nanopore DRS data, complementing established NGS-based structure-probing methods. Second, both segSHAPE and SMS-seq operate on the same input DRS data with the same DEPC probe, yet only segSHAPE captures the bound-state rearrangement, indicating that the signal-processing pipeline, rather than the chemistry, is the limiting factor for tracking conformational switches in this setting.

### 2.5 Structure prediction benchmark on RNA004 dataset

All preceding benchmarks were performed on RNA002 data. The recent RNA004 chemistry, however, uses a different pore model, a longer *k*-mer context (9-mer in RNA004 versus 5-mer in RNA002), and a different signal scale, so a pipeline tuned on RNA002 is not guaranteed to transfer to RNA004. To the best of our knowledge, the sm-PORE-cupine study [16] provides the only public RNA004 Nanopore DRS dataset currently available for SHAPE-based RNA structure probing. This study includes matched RNA002 and RNA004 data for the *Tetrahymena* ribozyme, together with an independent, non-nanopore structural reference from SAFA footprinting. Note that this *Tetrahymena* RNA002 dataset (NAI-N3 50 mM) is from [16] and is distinct from the *Tetrahymena* dataset (NAI-N3 100 mM) [13] benchmarked in Table 1. SAFA (Semi-Automated Footprinting Analysis) is a long-established tool that converts gel- and capillary-electrophoresis footprinting experiments into per-nucleotide reactivity values [33]; using the SAFA footprinting data reported by Wang et al. [16], we classified positions in the *Tetrahymena* ribozyme as single-stranded or double-stranded, providing a structural annotation independent of nanopore sequencing.

We applied segSHAPE and sm-PORE-cupine to both RNA002 and RNA004 datasets and evaluated their performance. We first assessed the quality of the per-position reactivity estimates against the SAFA-derived annotation. segSHAPE recovers single-stranded positions with per-position ROC-AUC values of 0.937 on RNA002 and 0.898 on RNA004, compared with 0.759 and 0.730 for sm-PORE-cupine on the same data (Figure 5A). Because single-stranded positions make up only 7% of the annotation, ROC-AUC can be mildly optimistic; we therefore also evaluated precision-recall performance, for which a no-skill classifier has a PR-AUC of 0.07. segSHAPE again leads decisively, with PR-AUC values of 0.558 on RNA002 and 0.538 on RNA004, compared with 0.298 and 0.284 for sm-PORE-cupine (Supplementary Figure S3). Thus, segSHAPE tracks the independent footprinting reference more closely than sm-PORE-cupine on both chemistries and under both metrics, and this advantage is retained as the chemistry changes from RNA002 to RNA004.

**Fig. 5.**
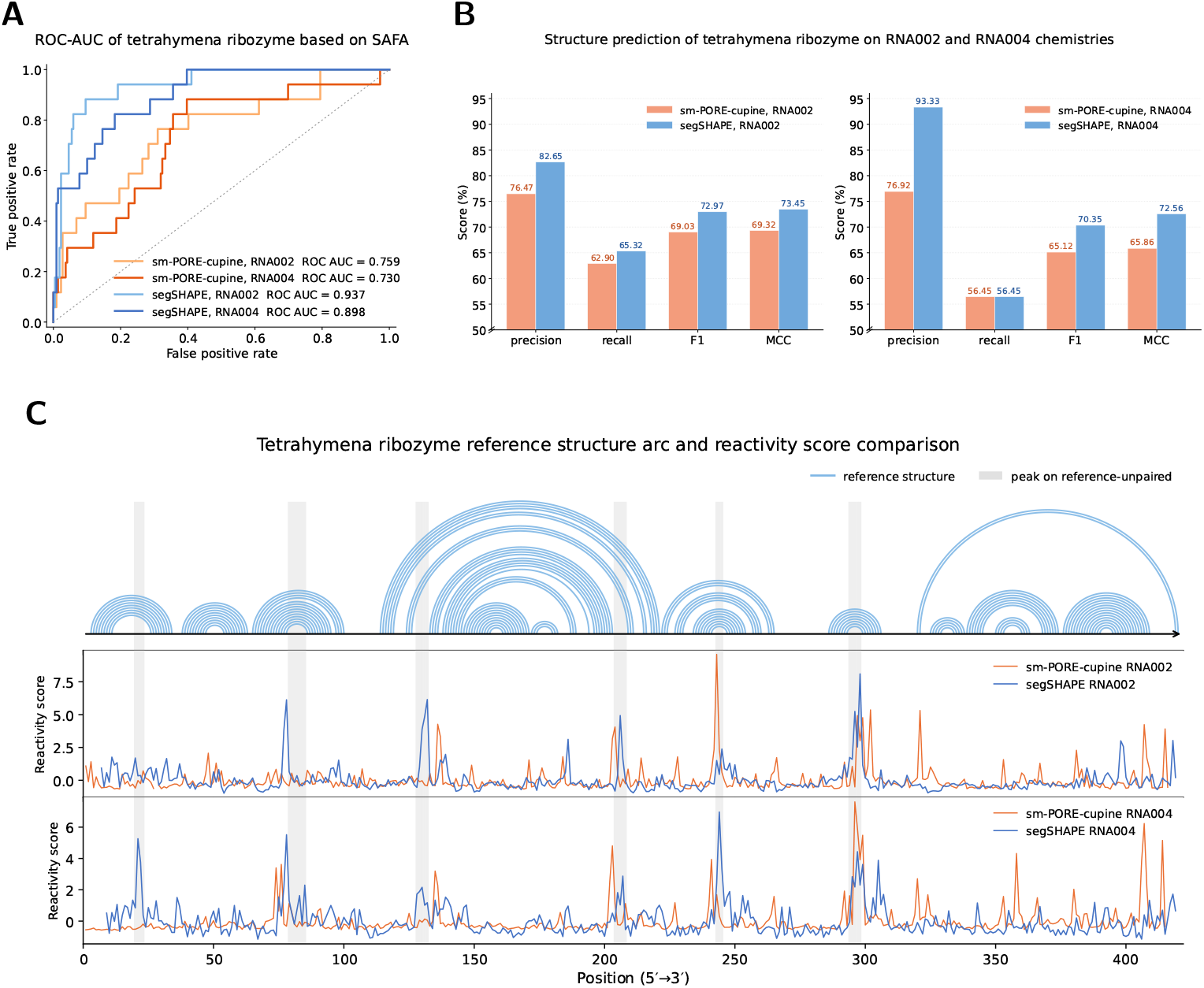
Comparison of segSHAPE and sm-PORE-cupine on *Tetrahymena* preribosomal RNA across RNA002 and RNA004 chemistries. (A) ROC curves evaluating per-position reactivity against the single-stranded/double-stranded annotation derived from SAFA footprinting of the *Tetrahymena* ribozyme RNA. (B) Structure-prediction metrics (precision, recall, F1 and MCC) on RNA002 and RNA004 chemistries, with the *y*-axis truncated at 50 to emphasize top-end differences. (C) Reference *Tetrahymena* secondary structure rendered as an arc diagram (top, 421 nt), aligned to the per-position reactivity profiles of segSHAPE (blue) and sm-PORE-cupine (orange) on RNA002 (middle) and RNA004 (bottom). Gray vertical bands mark high-reactivity peaks called by segSHAPE that fall in reference-unpaired regions, showing that segSHAPE consistently concentrates high reactivity in loop regions across both chemistries.

The structure-prediction metrics confirm this transferability (Figure 5B). On RNA002, segSHAPE outperforms sm-PORE-cupine on all four metrics, including a 4.1-point gain in MCC. On RNA004, segSHAPE retains higher precision, F1, and MCC, including a 16.4-point gain in precision and a 6.7-point gain in MCC. Direct inspection of the reactivity profiles shows that the high-reactivity peaks called by segSHAPE on both chemistries are concentrated in loop regions of the reference structure, with consistent peak locations across RNA002 and RNA004 (Figure 5C). These results indicate that the design choices in segSHAPE are not specific to the RNA002 pore model and transfer to the newer RNA004 chemistry, providing a forward-compatible pipeline as the community migrates away from RNA002.

## 3 Discussion

We presented segSHAPE, a unified computational framework for RNA secondary structure prediction from Nanopore DRS data. Across five public DRS benchmark datasets covering three chemical probes (AcIm, NAI-N3, DEPC) and RNAs from 165 to 1552 nt, segSHAPE achieves state-of-the-art accuracy on every benchmark RNA, including the in-vivo-probed *B. subtilis* 16S rRNA. It additionally captures ligand-induced riboswitch dynamics directly from DRS data and generalizes to the newer RNA004 chemistry.

The main conclusion from these results is that upstream signal interpretation is critical for DRS-guided structure prediction. Both segSHAPE variants share the same anchored alignment, per-read baseline correction and position-specific *k*-mer modeling, and differ only in the final per-position anomaly detector. Their similar and consistently strong performance suggests that the major gains come from cleaner signal alignment and a more accurate estimate of the unmodified per-position signal distribution, rather than from a particular outlier detector. This interpretation is consistent with the signal-distribution examples in Figure 2, where position-specific modeling resolves mismatched or multimodal control distributions that are not captured by a unified *k*-mer model.

A second observation is that supplying reactivity information does not automatically improve structure prediction. On two of the three benchmark RNAs (*Tetrahymena, B. subtilis* 16S rRNA), two reactivity-aware baselines (SMS-seq and nanoSHAPE) performed *worse* than reactivity-free RNAfold, indicating that low-quality reactivity profiles can actively harm prediction when upstream signal alignment is noisy. segSHAPE avoids this decline by improving the reactivity signal at its source, through position-specific *k*-mer modeling and basecalling-prior integration. These design choices also have broader implications for general Nanopore signal analysis beyond RNA structure probing [18].

The TPP riboswitch analysis illustrates the biological value of this improved signal processing. Unlike static benchmark structures, the TPP riboswitch requires recovery of a ligand-induced conformational rearrangement from two DRS experiments. segSHAPE reconstructs the expected ligand-free and TPP-bound topologies, whereas SMS-seq does not clearly resolve the switch. This result suggests that DRS-based reactivity profiles can support analysis of dynamic RNA structural changes when the upstream signal processing is sufficiently accurate.

The RNA004 comparison further separates two parts of the problem: detecting chemically perturbed positions from raw signal, and converting those reactivities into folding constraints. segSHAPE preserves high per-position ROC-AUC on RNA004, indicating that the signal-level ranking of modified or flexible positions transfers across chemistries. However, the lower RNA004 structure-prediction MCC suggests that the downstream SHAPE-to-energy conversion used by RNAfold may require chemistry-specific calibration. This will become increasingly important as the community moves from RNA002 to RNA004 and as additional DRS chemistries become available.

Deep learning-based RNA structure prediction methods, such as MXfold2 [30], provide a complementary route to structure prediction. Learned models exploit sequence-to-structure regularities from large training databases, whereas DRS-based reactivity pipelines inject experimental evidence for the specific RNA molecule being assayed. On our benchmark, MXfold2 achieves a competitive MCC on *Tetrahymena* (71.33), close to several reactivity-aware baselines, although its current implementation imposes a 1000-nt hard length limit that excludes long transcripts such as 16S rRNA. Combining learned sequence priors with DRS-derived reactivity constraints is therefore a natural direction for future structure-inference frameworks.

Three limitations of the present study should be noted. First, although our benchmark includes one in-vivo-probed RNA (*B. subtilis* 16S rRNA), the remaining datasets were probed *in vitro*; broader in-vivo benchmarking across more transcripts and cellular contexts is left for future work. Second, our RNA004 evaluation is currently limited to a single RNA (*Tetrahymena* ribozyme); broader RNA004 benchmarks await additional public structure-probing datasets with matched ground truth. Third, the anchored aligner can terminate prematurely on some reads, particularly in heavily modified or long transcripts. As a consequence, the fraction of reads passing the per-dataset coverage threshold (Supplementary Table S2) is consistently lower in treated than in matched control samples. Further refinement of the alignment to better handle such reads is left for future work.

## 4 Methods

### 4.1 Workflow overview

An overview of the segSHAPE workflow is presented in Figure 1. The pipeline takes raw current signal files (FAST5 or POD5) from matched control and chemically treated samples, together with a reference sequence, and returns the predicted RNA secondary structure. It comprises four main stages: (i) data preprocessing and signal segmentation for both samples; (ii) iterative anchored alignment to estimate a position-specific *k*-mer parameter table ***T*** from the control sample; (iii) alignment of the treated sample using the estimated ***T***, followed by per-position modification detection with Isolation Forest (default) or Gaussian mixture model; and (iv) reactivity calculation and SHAPE-guided structure prediction with RNAfold. The pipeline is implemented as a Python package with a single command-line entry point, `segshape,` which exposes each step as an independent sub-command (`pod5index, dorado-extract, segment, event-align, mod-calling, fold, evaluate, plot`); the package includes built-in *k*-mer tables for both RNA002 (5-mer) and RNA004 (9-mer) chemistries.

### 4.2 Data preprocessing

The pipeline accepts raw nanopore data in POD5 format. Legacy FAST5 inputs, which are typical of early RNA002 datasets, were first converted to POD5 using the official ONT `pod5` command-line toolkit, since current Dorado releases consume POD5 natively. All samples were then basecalled with Dorado in reference-guided mode, which internally calls minimap2 [34] to align the basecalled read sequence to the reference. RNA002 data were processed with Dorado v0.9.6 and RNA004 data with Dorado v1.4.0. In both cases, we used `--emit-moves` to produce BAM coordinates and a per-read `mv` (move) table. The resulting BAM files were filtered with samtools (v1.21) [35] using `-F 2324,` retaining only reads whose mapping coverage exceeded 80% of the reference length; this threshold is exposed as the `--min-coverage-fraction` flag and may be relaxed, for example to 50%, for datasets with shorter reads or sparser end-to-end coverage. For each read, we used `reference_start` and `reference_end` to define the aligned transcript interval on the reference, and the `ts`, `ns` and `mv` BAM tags to recover the corresponding raw-signal interval. This transcript-region signal extraction is taken directly from Dorado’s basecalling output and replaces the adapter/poly(A) trimming step used in earlier DRS pipelines, which depends on the Nanopolish `polya` module and is especially unreliable on short or degraded reads (Supplementary Figure S1). Finally, each read was robustly *z*-score-normalized using its own median and MAD: each raw-current value *s* was mapped to (*s* −median(*s*))*/*(1.4826· MAD(*s*)), where the factor 1.4826 makes the MAD an unbiased estimator of the Gaussian standard deviation. This per-read normalization removes pore-to-pore and run-to-run baseline drift and brings all reads onto a common scale.

### 4.3 Signal segmentation

The extracted normalized transcript-region signal is then segmented into sub-events using a lightweight find-peaks segmenter inspired by SegPore [17]. Specifically, we compute the absolute slope of the normalized signal, smooth this slope, and call `scipy.signal.find_peaks` with a fixed minimum peak-to-peak distance of 10 samples, with the detected peaks delimiting individual sub-events. For each sub-event, we calculate the mean and standard deviation of the in-segment current using a symmetric trimmed estimator: the leading and trailing 10% of samples in each segment are discarded before computing the mean and the standard deviation (exposed as `segshape segment --trim 0.1`). These trimmed regions correspond to transitions between adjacent nanopore states, where the current moves from one *k*-mer level to the next; including them would bias the mean toward neighboring levels and inflate the standard deviation through boundary jitter. Trimming therefore retains the plateau portion of each sub-event and reduces boundary-induced bias in the estimated subevent mean and standard deviation. This procedure produces a finer subdivision of the signal than the one-event-per-base representation used by Nanopolish, typically yielding 3∼ 7 sub-events per basecalled base. The resulting sub-events provide the downstream Viterbi dynamic programming algorithm with a denser set of indexable points and explicit entry and exit buffers around each basecalling-derived anchor. Representative find-peaks segmentations on four RNAs are shown in Supplementary Figure S4, and the full raw-signal normalization and segmentation procedure is given in Supplementary Algorithm S1.

### 4.4 Iterative alignment for position-specific parameter estimation

The untreated control sample is assumed to represent the unmodified signal distribution of the RNA. We use this sample to estimate a position-specific *k*-mer signal baseline at every reference position. A reference sequence of length *L* is represented as a sequence of overlapping *k*-mers, with *k* = 5 for RNA002 and *k* = 9 for RNA004; the *j*-th reference position denotes the *k*-mer centered at that position. The *k*-mer parameter tables distributed with segSHAPE use a common 5^*′*^→3^*′*^ canonical key convention, matching the convention used by Nanopolish and Dorado.

The position-specific parameter table defines a Gaussian baseline for each reference position:

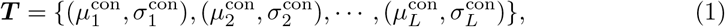

where 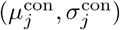 denote the mean and standard deviation of the unmodified signal distribution at reference position *j*. Given ***T***, each sub-event mean *e*_*u*_ is scored against position *j* using the Gaussian log-likelihood 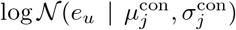. These per-event log-likelihoods are summed along an alignment path to obtain the read-level alignment score used by the dynamic-programming aligner described in the next section.

***T*** is initialized from the official ONT `kmer_models` repository (https://github.com/nanoporetech/kmer_models), supplemented with f5c (https://github.com/hasindu2008/f5c) [36]. For both chemistries, the per-position means 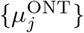 are taken directly from the normalized ONT level tables, which provide only normalized mean levels and no standard deviations; per *k*-mer standard deviations are available only in picoampere (pA) units. To place the standard deviations on the same normalized scale as the means, we regressed the normalized level means (*µ*^z^) against the corresponding pA-scale means 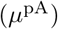 across all *k*-mers, 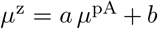. We empirically obtained the normalized standard deviations as 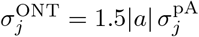. Specifically, for RNA002 the means are taken from `rna_r9.4_180mv_70bps/5mer_levels_v1.txt` and the pA-scale standard deviations from the `legacy legacy_r9.4_180mv_70bps_5mer_RNA/template_median69pA.model` table, giving *a* = 0.05401 (*b* = − 4.973, *r*^2^ = 1.000). For RNA004 the means are taken from `rna004/9mer_levels_v1.txt` and the pA-scale standard deviations from the `model_pre_r10`9-mer template distributed with f5c (file `src/model.h; SHA acb3826`), giving *a* = 0.05728 (*b* = − 4.827, *r*^2^ = 0.996). These chemistry-specific tables are loaded once at startup and used as the ONT prior 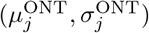 throughout iterative refinement.

***T*** is estimated by alternating between alignment and parameter update steps. In the alignment step, each control read is aligned to the reference under the current ***T*** using the anchored aligner described in the next section. In the parameter update step, the mean at each position *j* is re-estimated as a Bayesian shrinkage estimate that combines the empirical mean of all sub-event means currently aligned to *j* with the ONT *k*-mer prior 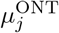 with prior weight *κ*:

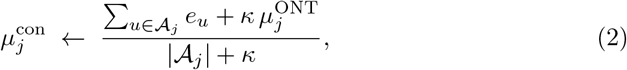

where *A*_*j*_ is the set of control sub-events currently aligned to position *j*. The prior weight *κ* = 50 (in units of effective sub-events) keeps low-coverage positions anchored to the global ONT baseline and prevents them from drifting on a handful of stray sub-events. ***T*** is iterated for *Q* = 3 outer rounds, after which the converged ***T*** is frozen and used as input to the treated-sample alignment. Pseudocode of the position-specific parameter estimation is provided in Supplementary Algorithm S4.

### 4.5 Anchored alignment and per-read baseline shift correction

Alignment of signal sub-events to the reference sequence (steps 3 and 5 of the pipeline) is constrained by basecalling-derived anchor points and complemented by a per-read baseline shift correction. Together, these two components address two common failure modes of unconstrained DRS signal alignment: phantom matching due to mis-specified transcript region boundaries in the raw signals, and low alignment scores due to per-read baseline drift. Supplementary Figure S2 provides an illustration of the anchored alignment algorithm, and the complete pseudocode is given in Supplementary Algorithms S2 to S4.

#### Anchor points from Dorado’s move table

Dorado emits a move table (the `mv` BAM tag) that records, for each block of raw-signal samples consumed by the neural decoder, whether the decoder advances to a new base; cumulatively summing these moves yields a coarse but robust signal-to-base correspondence directly from basecalling. In reference-guided mode, Dorado produces BAM records that align each basecalled read to the reference, allowing the contiguous reference-matching region to be read directly from the BAM record; we then recover the corresponding signal-sample indices from `mv`. The resulting (start, end) pairs, expressed both as reference *k*-mer indices (*k*_seed_, *k*_end_) and as sub-event indices (*j*_min_, *j*_max_), are the per-read *anchors* (Supplementary Figure S2).

#### Box-constrained Viterbi dynamic programming (DP) algorithm

Because the basecalling anchors are approximate, we relax each anchor by *δ*_*k*_ reference *k*-mers and *δ*_*e*_ sub-events in both directions, defining a per-read *entry box* around (*k*_seed_, *j*_min_) and an *exit box* around (*k*_end_, *j*_max_ + 1) (Supplementary Figure S2). For sub-event index *u* ∈{1, …, *S*} and reference *k*-mer index *j* ∈{1, …, *L* }, we run a Viterbi DP algorithm with the recurrence

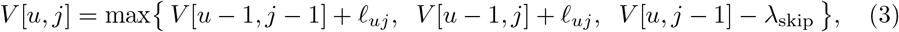

corresponding respectively to a match, a stay (multiple sub-events sharing one *k*-mer) and a *k*-mer skip with constant penalty *λ*_skip_. The per-cell match score is

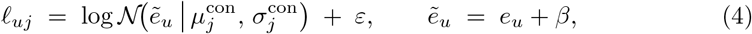

where 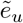 is the sub-event mean after a per-read baseline shift *β* (introduced below), and *ε* is a per-event offset that calibrates match and skip on the same scale, set automatically to 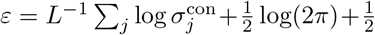 (see Supplementary Algorithm S3 and Supplementary Algorithm S4). The DP is initialized from every cell in the entry box, propagated through (3), and the highest-scoring exit-box endpoint is taken as the optimal alignment. Parameter defaults are *δ*_*e*_ = 50, *δ*_*k*_ = 15, and *λ*_skip_ = 50. The full procedure is detailed in Supplementary Algorithm S2.

#### Per-read baseline shift correction

Within a single alignment, *β* is refined by alternating the box-constrained Viterbi DP with a robust re-estimate from the current alignment,

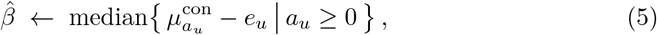

where *a*_*u*_ is the reference position to which sub-event *u* is currently aligned and *a*_*u*_ ≥0 excludes unaligned sub-events. Using the median rather than the mean makes (5) robust to the small fraction of outlier positions arising from chemical modifications, stalls or transient blockages. Per-read iteration uses up to *R* = 3 inner rounds with log-likelihood tolerance 10^−1^ for early stopping, and 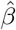 is clipped to a chemistry-dependent envelope to prevent pathological drift on poorly aligned reads ([−1, 1] for RNA002, [− 0.3, 0.3] for RNA004). The same per-read aligner is reused inside the outer iteration that refines the position-specific parameter table ***T***, whose full schedule is given in Supplementary Algorithm S4.

### 4.6 Modification estimation using Isolation Forest

Normalized signal sub-events from the treated sample are first aligned to the reference sequence using the parameter table ***T*** estimated from the control sample. For each reference position *i*, we collect the control sub-event means aligned to that position and use Isolation Forest [23] to model the unmodified signal distribution. Treated-sample sub-events that deviate from this position-specific control distribution are then flagged as putative chemical modifications.

Let ***X*** ={ ***x***_1_, …, ***x***_*L*_}denote the final-iteration control alignment, where ***x***_*i*_ is the set of sub-event means aligned to reference position *i* and *L* is the reference length. For each position *i*, an Isolation Forest with *M* trees is constructed from ***x***_*i*_ by recursively partitioning the control sub-event means at random split values until each leaf isolates a single point or a maximum depth is reached. For a query *x*, the anomaly score follows the standard Isolation Forest formulation [23]:

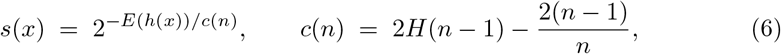

where *E*(*h*(*x*)) is the average path length of *x* across the forest, *n* = |***x***_*i*_|, and 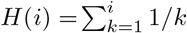. The normalization *c*(*n*) is the expected path length of an unsuccessful search in a random binary search tree, which mirrors the recursive partitioning of an isolation tree [37]. Higher *s*(*x*) values indicate stronger deviation from the control distribution.

Let ***y***_*i*_ denote the set of treated-sample sub-event means aligned to position *i*. Applying the corresponding per-position forest to ***y***_*i*_ yields anomaly scores *{s*(*y*_*ij*_)*}*_*j*_; the modification rate *α*_*i*_ is defined as the fraction of treated sub-events whose score exceeds the threshold set by the `contamination`parameter. The full modification-rate vector is ***α*** = (*α*_1_, …, *α*_*L*_). In practice, we use scikit-learn (v1.3.0) [38] `IsolationForest` with `contamination=0.005,` corresponding to a 0.5% expected outlier rate in the control sample and matching the same GMM cut-off.

### 4.7 Modification estimation using Gaussian mixture model

As an alternative to Isolation Forest, segSHAPE provides a Gaussian mixture model (GMM) outlier detector; this variant is reported as segSHAPE (GMM) in Table 1. The motivation is that, after the iterative position-specific parameter table ***T*** is estimated, the per-position control distribution ***x***_*i*_ is expected to be close to a unimodal Gaussian centered on 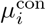, whereas chemically modified treated reads can introduce a second component with a shifted mean. A low-component GMM therefore provides a simple label-free model of the unmodified control distribution while allowing residual substructure when supported by the data.

For each reference position *i*, we fit a GMM to ***x***_*i*_ using scikit-learn’s (v1.3.0) `GaussianMixture`class [38]. The number of mixture components *K*_*i*_ ∈{1, 2} is selected per position by the Bayesian information criterion (BIC): both candidates are fitted when |***x***_*i*_| ≥20, otherwise only *K*_*i*_ = 1 is considered, and the model with the lower BIC is retained. This selection procedure reduces to a single Gaussian when the control signal is unimodal or insufficiently sampled, while allowing a second component when residual subpopulations are supported by the data.

Let *p*_*i*_(·) denote the resulting log-probability under the selected GMM. We define the per-position cut-off *τ*_*i*_ as the *q*-th quantile of the control log-probabilities {*p*_*i*_(*x*_*ij*_)}_*j*_ with *q* = 0.005, matching the expected control-sample outlier rate of 0.5% used by the Isolation Forest outlier detector. This matched calibration makes the two outlier detectors directly comparable in Table 1 and Supplementary Table S3. The modification rate at position *i* is then the fraction of treated log-probabilities below *τ*_*i*_,

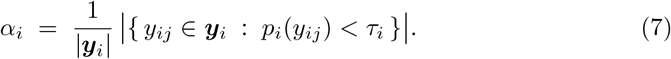

Both outlier detectors use the same per-position aligned events and differ only in the final modification-detection step; the rationale for adopting Isolation Forest as the default is given. Because their tail thresholds are calibrated to the same 0.5% control-outlier rate, the two detectors remain directly comparable: Isolation Forest gives the higher F1 and MCC on pri-miR-17 ∼ 92 and *Tetrahymena*, whereas the GMM performs better on *B. subtilis* 16S rRNA. The full per-dataset comparison is provided in Supplementary Table S3.

### 4.8 Reactivity normalization and structure folding

The per-position modification rates ***α*** are first *z*-score normalized to place modification rates from different chemical probes on a common dynamic range. The resulting normalized reactivity vector ***β*** is written as a SHAPE `.dat` file. Positional smoothing is exposed as a configurable parameter (`--smooth-window`) and is set to 0 by default, corresponding to no smoothing. The reactivity vector and reference sequence are then provided to RNAfold (v2.7.0) [21], using the options `-p -d2 --noLP -P rna_andronescu2007.par --shapeMethod=D.` RNAfold converts the supplied reactivities into SHAPE-derived pseudo-energies during folding, and the resulting centroid structure is taken as the final segSHAPE output. For visualization, base-pairing patterns of centroid structures are rendered as arc diagrams using R-chie (v0.1.3) [39].

### 4.9 Benchmarking and ablation study

For benchmarking on RNA002 data, we selected six baseline methods. MXfold2 [30], RNAPKplex (v2.7.0), and RNAfold (v2.7.0) [21] predict RNA secondary structures from the reference sequence alone, whereas SMS-seq [12], nanoSHAPE [11], and PORE-cupine [13] additionally use experimentally derived reactivity scores. The evaluation datasets comprise pri-miR-17 ∼ 92 RNA [11], *Tetrahymena* RNA, and *Bacillus subtilis* 16S rRNA [13] for the main RNA002 benchmark; the thiamine pyrophosphate (TPP) riboswitch [12] for ligand-dependent structural rearrangement; and the matched *Tetrahymena* RNA004 dataset [16] for evaluating generalization to RNA004 chemistry. All raw nanopore data were retrieved from public repositories, with accessions and per-sample statistics listed in Supplementary Tables S1 and S2. For the RNA004 comparison, per-position modification rates for sm-PORE-cupine were taken from the original publication [16]. The upstream segSHAPE pipeline was run with identical default parameters on all datasets; only the two downstream reactivity-processing parameters for TPP riboswitch and RNA004 datasets were re-calibrated. Full parameter settings are given in Supplementary Methods 4.1–4.2.

To isolate the contribution of the per-position outlier detector from the rest of the pipeline, we compared the segSHAPE default (Isolation Forest) and the GMM variant with five alternative modification-detection strategies while holding the upstream pipeline fixed. The seven methods fall into two families. *Anomaly-detection* methods model only the control distribution and flag deviations in the treated sample; this group includes Isolation Forest, GMM with a tail-quantile cut-off, One-Class SVM, |Δmedian|, the Kolmogorov-Smirnov two-sample statistic, and the 1-D Wasserstein distance. *Differential-mixture* methods jointly model the control and treated samples and return per-position modification rates directly; this family is represented by xPore [28]. For a fair comparison, methods with a tail-fraction hyperparameter were calibrated to the same 0.5% expected control-sample outlier rate; the remaining methods (|Δmedian |, KS, Wasserstein, and xPore) were used without additional tuning. Each variant was passed through the same downstream pipeline: per-position *z*-score normalization, smoothing window = 0, and RNAfold centroid prediction. All variants were evaluated against the reference structure using precision, recall, F1, and MCC, with full per-dataset results reported in Supplementary Table S3. Further benchmarking details are provided in Supplementary Methods 4.3.

### 4.10 Reference structures of RNAs

The reference structures for *Tetrahymena* RNA and *Bacillus subtilis* 16S rRNA were obtained from the Comparative RNA Web (CRW) database [40]. We excluded pseudoknot base pairs for compatibility with RNAfold-based prediction [21, 41], yielding 124 and 470 nested reference base pairs for *Tetrahymena* and 16S rRNA, respectively. The structure of the TPP riboswitch was obtained from a previous study [42]. The reference structure for pri-miR-17∼ 92 was obtained by RNAfold prediction guided by published SHAPE-MaP reactivity data [11].

### 4.11 Structure prediction evaluation

We evaluated the predicted RNA secondary structures against reference (ground truth) structures using four standard metrics: precision (P, also known as positive predictive value), recall (R, also referred to as sensitivity or true positive rate), the F1 score, and the Matthews correlation coefficient (MCC) [21, 43, 44]. All four are based on the binary classification of nucleotide positions as paired or unpaired, derived from true positives (TP), true negatives (TN), false positives (FP), and false negatives (FN). We report all four for completeness but treat MCC as the primary metric, as it jointly accounts for all four confusion-matrix categories and remains reliable under the paired/unpaired class imbalance intrinsic to base-pair classification [31]. For brevity, we use the abbreviations P and R in Table 1 and in the supplementary tables. An illustration of a hairpin structure together with a toy example of base-pair-level TP/FN/FP/TN counting is provided in Supplementary Figure S5.

### 4.12 SAFA-based evaluation

In addition to the base-pair-level metrics above, we evaluated per-position reactivity on the *Tetrahymena* ribozyme (Figure 5A) against an independent, nanopore-free reference: the SAFA footprinting data from Extended Data Fig. 6i of Wang et al. [16], which provides a single-stranded versus double-stranded annotation over reference positions 121 to 356. Treating the 17 single-stranded positions as the positive class and the 219 double-stranded positions as the negative class, we computed the ROC curve and its area (ROC-AUC) using each method’s per-position reactivity as the classifier score. Because single-stranded positions constitute only 17 of the 236 annotated positions, we additionally report the precision-recall curve and its area (PR-AUC), which does not depend on the large negative class and is therefore less sensitive to this imbalance (Supplementary Figure S3). The same SAFA-derived annotation was applied to both the RNA002 and RNA004 data and to both segSHAPE and sm-PORE-cupine, so that the comparison in Figure 5A is fully matched.

## 5 Data and code availability

All raw FAST5 or POD5 data used in this study were obtained from public repositories. The pri-miR-17 ∼ 92 RNA FAST5 data are available from NCBI SRA under accession number PRJNA634693. The *Tetrahymena* RNA (RNA002, NAI-N3 100 mM) and *Bacillus subtilis* 16S rRNA FAST5 data are available from Gene Expression Omnibus (GEO) under accession number GSE133361. The *Tetrahymena* RNA (RNA002, NAI-N3 50 mM) is available under accession number GSE304702, and the matched *Tetrahymena* RNA004 dataset under accession number GSE304703. The TPP riboswitch data are available from the European Nucleotide Archive (ENA) under accession code PRJEB36658. Further details of all datasets are provided in Supplementary Tables S1 and S2.

segSHAPE is distributed as a Python package with a single command-line entry point (`segshape`) that exposes each pipeline step as an independent sub-command (`pod5index / dorado-extract / segment / event-align / mod-calling / fold / evaluate / plot`); all result data, source code and example workflows are available on GitHub (https://github.com/guangzhaocs/segSHAPE).

## 6 Supplementary data

Supplementary data is available alongside this preprint.

## 7 Competing interests

No competing interest is declared.

## 8 Author contributions statement

G.C. and L.C. conceived the study. G.C. performed the majority of the experiments and analyses, with L.H. contributing to per-read signal baseline shift correction idea and experiments. M.-L.A. provided advice and suggestions. G.C. and L.C. wrote the manuscript. L.C. supervised the project. All authors reviewed and approved the final manuscript.

## 9 Acknowledgments

We acknowledge the computational resources provided by the Aalto Science-IT project, CSC – IT Center for Science, Finland, and UEF Bioinformatics Center, Biocenter Kuopio, Biocenter Finland, University of Eastern Finland, Finland. G.C. and L.C. thank the Research Council of Finland grants (No. 335858, 358086). G.C. thanks the Finnish Cultural Foundation grant.

## Notes

### Competing Interest Statement

The authors have declared no competing interest.

