## Supplementary Materials for "segSHAPE: RNA secondary structure prediction from nanopore direct RNA sequencing"

### segSHAPE Supplementary Materials

June 2026

**References** **17**

### 1 Supplementary Figures

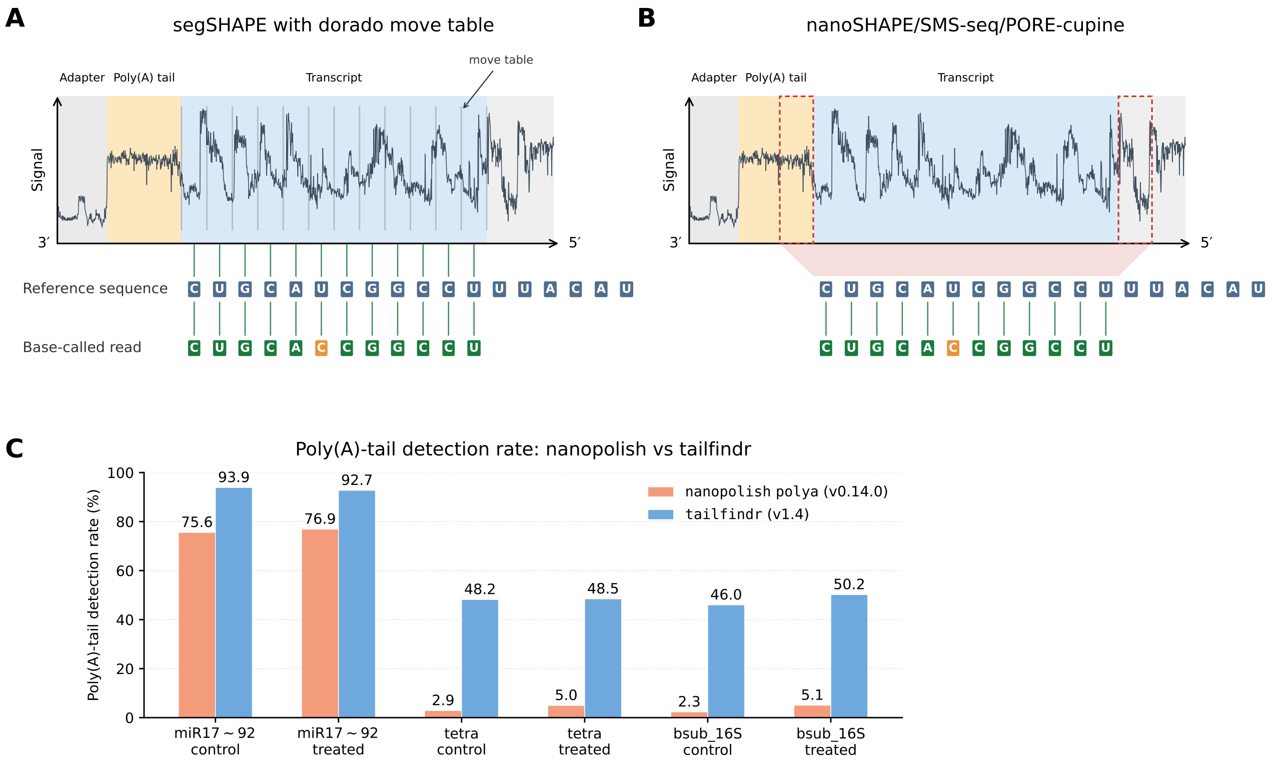

**Supplementary Figure S1: (A, B)** Schematic illustration of how the transcript region is delimited before signal-to-reference alignment. The raw current is shown $3' \to 5'$ with the adapter (gray), poly(A) tail (orange) and transcript (blue) regions indicated. The base-called-read-to-reference correspondence is identical in both panels; the panels differ only in where the boundary between poly(A)/adapter and transcript is placed upstream. **(A)** segSHAPE delimits the transcript region directly from Dorado's `ts`/`ns` tags and the `move` table, placing this boundary at the true junction. **(B)** Pipelines that rely on poly(A)/adapter trimming misplace the junction (red dashed boxes), so part of the poly(A) tail or a truncated $5'$ end is included in the transcript region and carried into the signal-to-reference alignment (phantom matching), contaminating or truncating the extracted transcript signal. **(C)** Poly(A)-tail detection rate of nanopolish `polya` (v0.14.0) [1] versus tailfindr (v1.4) across the three RNA002 benchmark datasets (pri-miR-17∼92, *Tetrahymena*, *B. subtilis* 16S rRNA), shown separately for the control and treated samples. The detection rate is the fraction of basecalled reads for which each tool returned a usable poly(A) call (`qc_tag = PASS` for nanopolish; a non-missing `tail_length` for tailfindr), taken over all reads in that tool's output.

Anchored Viterbi DP alignment: entry / exit boxes and backtrace path

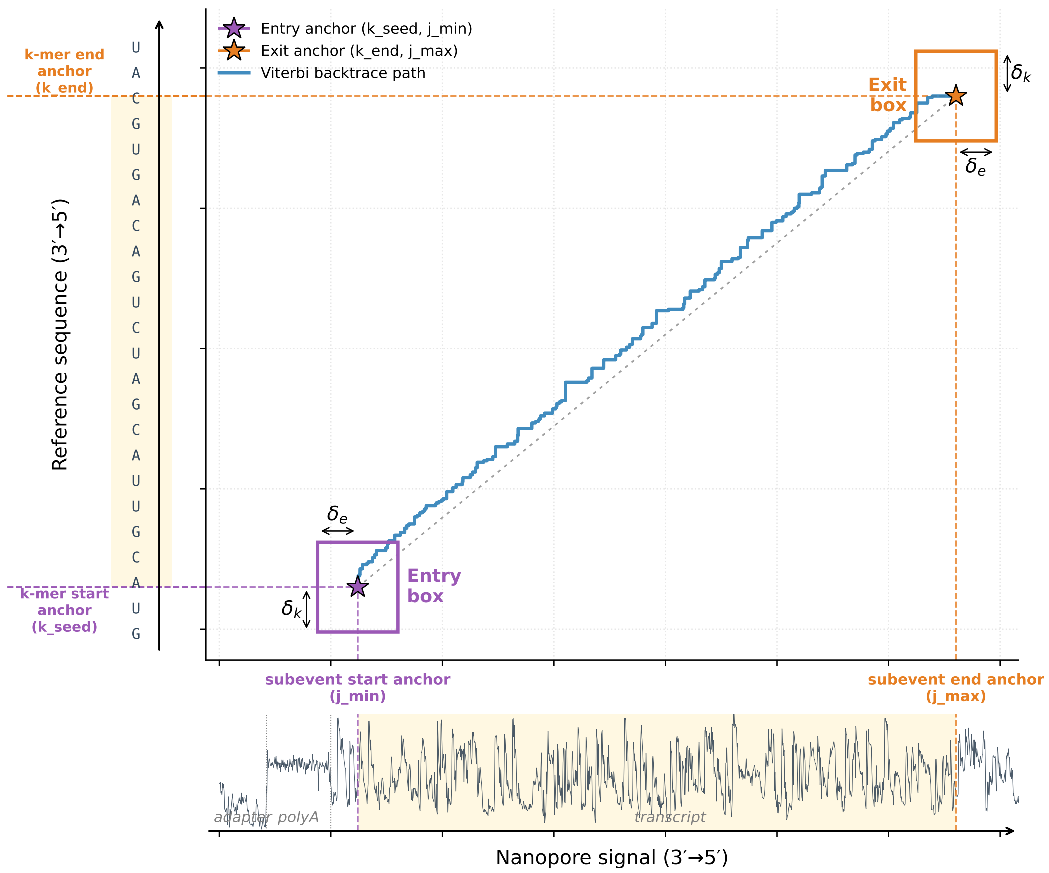

**Supplementary Figure S2:** Detailed view of the anchored alignment between the reference sequence (vertical axis, $k$-mer index) and the raw signal sub-events (horizontal axis, sub-event index). Basecalling-derived anchor points define an entry box around the start anchor $(k_{\text{seed}}, j_{\text{min}})$ and an exit box around the end anchor $(k_{\text{end}}, j_{\text{max}} + 1)$, each relaxed by $\delta_k = 15$ $k$-mers and $\delta_e = 50$ sub-events. The box-constrained Viterbi DP enumerates all admissible (start, end) combinations and returns the best-scoring alignment path. Each colored curve corresponds to one such candidate combination; the bold path is the selected best alignment.

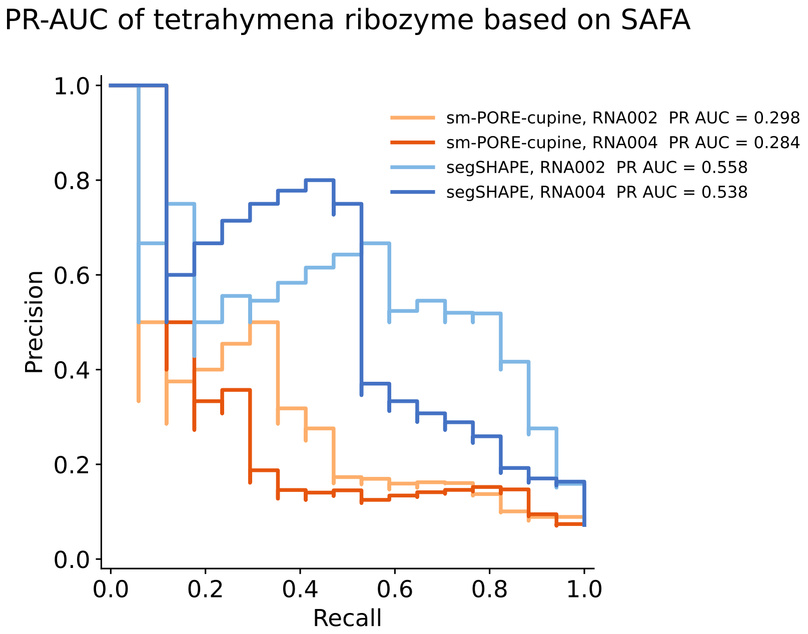

**Supplementary Figure S3:** Precision–recall (PR) curves of per-position single-strand accessibility on the *Tetrahymena* ribozyme using SAFA as the binary reference. Four traces compare the sm-PORE-cupine baseline against segSHAPE on both RNA002 (5-mer) and RNA004 (9-mer) chemistries, computed from the per-position raw modification rate (baseline) or the segSHAPE reactivity $z$-score (ours). The legend reports the area under the PR curve (PR-AUC) for each method/chemistry combination. segSHAPE improves PR-AUC over the matched sm-PORE-cupine baseline on both chemistries.

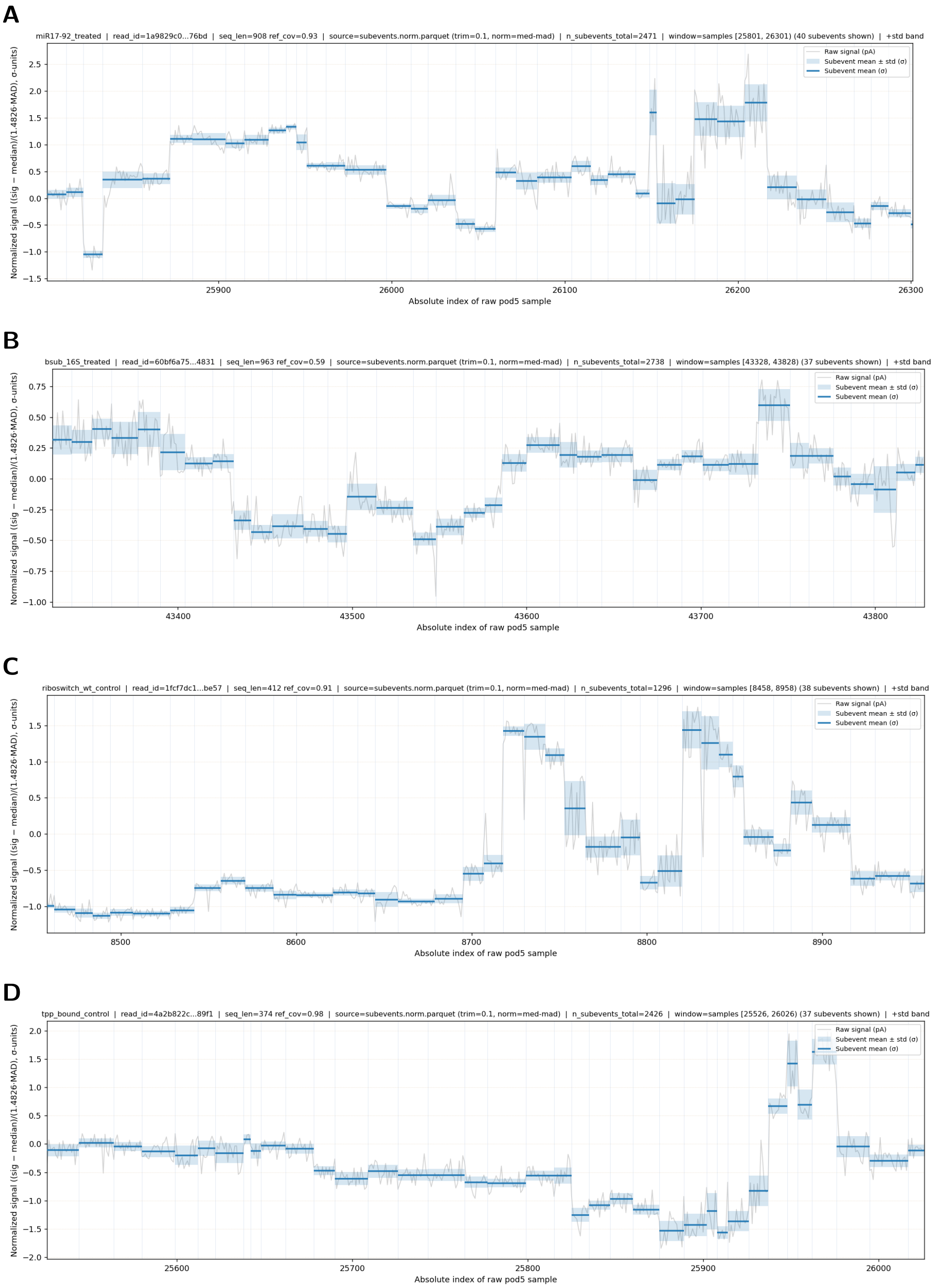

**Supplementary Figure S4:** Examples of slope-based find-peaks segmentation on four representative reads from different datasets. In each panel, the signal trace shows the per-read $z$-score normalized raw signal in a 500-sample window. Vertical dashed lines mark sub-event boundaries, and within each sub-event the blue line and the surrounding shaded band indicate the in-segment mean and standard deviation, respectively, computed after 10 % trimming from each sub-event. (A) Example read from the miR17∼92 treated sample. (B) Example read from the *B. subtilis* 16S rRNA treated sample. (C) Example read from the Tetrahymena RNA004 control sample. (D) Example read from the TPP-bound control sample. The segmenter consistently places boundaries at clear current-level transitions on both RNA002 (A, B, D) and RNA004 (C) chemistries.

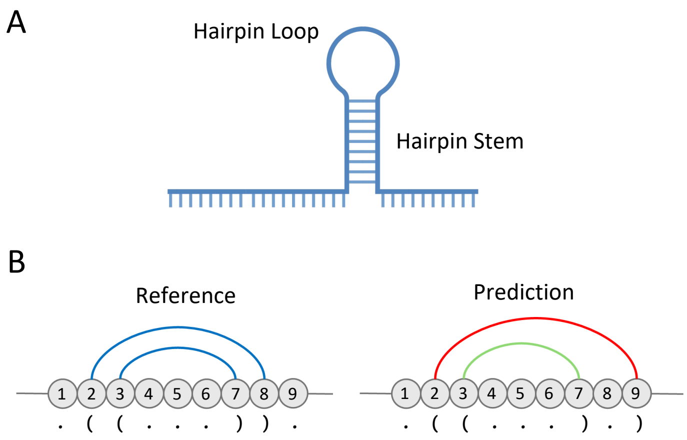

**Supplementary Figure S5:** Illustration of hairpin structure and evaluation metrics. (A) Schematic of a hairpin loop, with paired bases forming the stem regions and unpaired bases forming the loop. (B) Toy example of base-pair-level evaluation. The reference structure contains paired bases at positions $(2, 8)$ and $(3, 7)$, while the predicted structure contains base pairs $(2, 9)$ and $(3, 7)$. The pair $(3, 7)$ is classified as a true positive (TP), $(2, 8)$ as a false negative (FN), and $(2, 9)$ as a false positive (FP). True negatives (TN) are all remaining base pairs $(i, j)$ with $i, j \in [1, 9]$ and $i < j$ that appear in neither structure. With a sequence length of 9 there are $\binom{9}{2} = 36$ candidate pairs in total, giving $\mathrm{TP} = 1$, $\mathrm{FN} = 1$, $\mathrm{FP} = 1$, and $\mathrm{TN} = 36 - 3 = 33$. The four standard metrics then evaluate to $\mathrm{Precision} = \mathrm{TP}/(\mathrm{TP}+\mathrm{FP}) = 1/2 = 0.500$, $\mathrm{Recall} = \mathrm{TP}/(\mathrm{TP}+\mathrm{FN}) = 1/2 = 0.500$, $\mathrm{F}_1 = 2\,\mathrm{TP}/(2\,\mathrm{TP}+\mathrm{FP}+\mathrm{FN}) = 2/4 = 0.500$, and $\mathrm{MCC} = (\mathrm{TP}\cdot\mathrm{TN} - \mathrm{FP}\cdot\mathrm{FN})/\sqrt{(\mathrm{TP}+\mathrm{FP})(\mathrm{TP}+\mathrm{FN})(\mathrm{TN}+\mathrm{FP})(\mathrm{TN}+\mathrm{FN})} = (1\cdot 33 - 1\cdot 1)/\sqrt{2\cdot 2\cdot 34\cdot 34} = 32/68 = 8/17 \approx 0.471$.

#### 2 Supplementary Tables

**Supplementary Table S1**: Dataset experimental metadata. For each dataset we list the source publication, the SHAPE/acylation probe applied to the treated sample, its concentration, and whether probing was performed *in vitro* or *in vivo*. Each dataset has a matched untreated control (not listed). Per-sample sequencing and mapping statistics for the same datasets are given in Supplementary Table S2.

| dataset | source | chemistry | probe | concentration | condition |
|---|---|---|---|---|---|
| pri-miR-17∼92 | Stephenson *et al.*, 2022 [2] | RNA002 | AcIm | 150 mM | *in vitro* |
| *Tetrahymena* | Aw *et al.*, 2021 [3] | RNA002 | NAI-N3 | 100 mM | *in vitro* |
| *B. subtilis* 16S | Aw *et al.*, 2021 [3] | RNA002 | NAI-N3 | 100 mM | ***in vivo*** |
| TPP (bound) | Bizuayehu *et al.*, 2022 [4] | RNA002 | DEPC | 10% (v/v) | *in vitro* |
| TPP (unbound) | Bizuayehu *et al.*, 2022 [4] | RNA002 | DEPC | 10% (v/v) | *in vitro* |
| smPC_002pool_wt | Wang *et al.*, 2026 [5] | RNA002 | NAI-N3 | 50 mM | *in vitro* |
| smPC_004pool_wt | Wang *et al.*, 2026 [5] | RNA004 | NAI-N3 | 50 mM | *in vitro* |

**Supplementary Table S2**: Dataset statistics. In this table, chemistry is the Nanopore RNA kit (RNA002 = 5-mer pore model, RNA004 = 9-mer pore model); #pod5 is the number of input pod5 files; #reads is the number of reads in those pod5 files; min-ref-coverage is the minimum reference coverage applied in `segshape dorado-extract` (default 0.8, loosened to 0.5 on short or truncated transcripts); #mapped is the number of input reads that survived all `dorado-extract` filters (primary mapping, MAPQ $\geq$ 20, coverage cutoff, duplicate removal, and contig keep-list for riboswitch); accession lists the public archive entries from which the raw data were retrieved; the experimental metadata for each dataset (source, probe, concentration, and in vitro/in vivo condition) is summarized in Supplementary Table S1. For smPC_002pool_wt and smPC_004pool_wt, all samples were filtered with `--keep-contig TETRA`, so only TETRA-mapped reads contribute to their #mapped column.

| dataset | sample | accession | chemistry | #pod5 | #reads | min-ref-cov | #mapped |
|---|---|---|---|---|---|---|---|
| pri-miR-17∼92 | control | SRX8387629 | RNA002 | 69 | 68,693 | 0.80 | 12,849 |
| pri-miR-17∼92 | treated | SRX8387629 | RNA002 | 78 | 77,992 | 0.80 | 7,698 |
| *Tetrahymena* | control | GSM4490611 | RNA002 | 12 | 40,581 | 0.80 | 27,912 |
| *Tetrahymena* | treated | GSM4490616 | RNA002 | 19 | 72,099 | 0.80 | 41,328 |
| bsub_16S | control | GSM4490617 | RNA002 | 6 | 20,678 | 0.50 | 8,909 |
| bsub_16S | treated | GSM4490618 | RNA002 | 24 | 92,576 | 0.50 | 28,641 |
| TPP (bound) | control | PRJEB36658 | RNA002 | 6 | 21,409 | 0.50 | 6,653 |
| TPP (bound) | treated | PRJEB36658 | RNA002 | 11 | 41,614 | 0.50 | 5,765 |
| TPP (unbound) | control | PRJEB36658 | RNA002 | 19 | 74,149 | 0.50 | 26,779 |
| TPP (unbound) | treated | PRJEB36658 | RNA002 | 27 | 104,641 | 0.50 | 16,041 |
| smPC_002pool_wt | control | SRR36194983 | RNA002 | 1,061 | 4,243,270 | 0.80 | 45,012 |
| smPC_002pool_wt | treated | SRR36194985 | RNA002 | 306 | 1,221,342 | 0.80 | 11,121 |
| smPC_004pool_wt | control | SRR36194786 | RNA004 | 609 | 2,430,396 | 0.80 | 28,587 |
| smPC_004pool_wt | treated | SRR36194785 | RNA004 | 992 | 3,963,420 | 0.80 | 34,609 |

**Supplementary Table S3**: Ablation study of per-position modification-scoring methods on the three RNA002 benchmark datasets (pri-miR-17∼92, *Tetrahymena* preribosomal RNA, *B. subtilis* 16S rRNA). **All seven methods consume the same per-position events produced by the segSHAPE upstream pipeline** (anchored alignment with basecalling prior, position-specific $k$-mer model, and per-read baseline shift correction); only the per-position scoring step is swapped. This is therefore an ablation of the modification detector, not a comparison of end-to-end pipelines. The methods fall into two families: (i) unsupervised *anomaly-detection* detectors that model the control distribution and flag deviations in the treated sample (IF, GMM with tail-quantile threshold, OC-SVM, dmed, KS, Wasserstein); and (ii) unsupervised *differential-mixture* detectors that jointly model control and treated data with a two-component mixture and report per-position modification rate (xPore). To make the comparison fair, all detectors that expose a tail-fraction hyper-parameter were calibrated to the same expected control-sample outlier rate $\sim 0.5\%$: `contamination = 0.005` for IF, `nu = 0.005` for OC-SVM, and the 0.5% quantile of the control GMM log-probability for GMM. The remaining detectors (dmed, KS, Wasserstein, and xPore) are parameter-free. Reported metrics are precision (P), recall (R), F1 and MCC (all in %) computed on the RNAfold centroid structure. Within each dataset, baselines are listed first (ordered by MCC, descending) and the two model-based segSHAPE variants are placed at the bottom. Best value per column is shown in **bold**.

| Method | Precision(P) | Recall(R) | F1 | MCC |
|---|---|---|---|---|
| **pri-miR-17∼92** (RNA002, 951 nt, AcIm) | | | | |
| Wasserstein | 87.50 | 77.48 | 82.19 | 82.33 |
| xPore | **95.29** | 61.83 | 75.00 | 76.75 |
| dmed | 82.21 | 65.27 | 72.77 | 73.24 |
| KS | 71.11 | 48.85 | 57.92 | 58.92 |
| OC-SVM | 64.55 | 46.56 | 54.10 | 54.80 |
| segSHAPE (GMM) | 85.71 | 82.44 | 84.05 | 84.05 |
| segSHAPE (IF) | 89.84 | **84.35** | **87.01** | **87.04** |
| ***Tetrahymena* preribosomal RNA** (RNA002, 421 nt, NAI-N3) | | | | |
| xPore | **88.46** | 55.65 | 68.32 | 70.13 |
| KS | 67.90 | 44.35 | 53.66 | 54.83 |
| OC-SVM | 62.79 | 43.55 | 51.43 | 52.24 |
| Wasserstein | 50.45 | 45.16 | 47.66 | 47.66 |
| dmed | 50.47 | 43.55 | 46.75 | 46.81 |
| segSHAPE (GMM) | 83.50 | 69.35 | 75.77 | 76.07 |
| segSHAPE (IF) | 83.02 | **70.97** | **76.52** | **76.73** |
| ***B. subtilis* 16S rRNA** (RNA002, 1552 nt, NAI-N3) | | | | |
| xPore | 66.56 | 44.04 | 53.01 | 54.13 |
| Wasserstein | 58.91 | 41.49 | 48.69 | 49.42 |
| OC-SVM | 64.55 | 36.81 | 46.88 | 48.73 |
| dmed | 51.37 | 27.87 | 36.14 | 37.82 |
| KS | 47.74 | 20.21 | 28.40 | 31.05 |
| segSHAPE (GMM) | **70.44** | **54.26** | **61.30** | **61.81** |
| segSHAPE (IF) | 65.12 | 50.85 | 57.11 | 57.53 |

### 3 Supplementary Pseudo-Codes

**Supplementary Algorithm S1:** Slope-based find-peaks signal segmentation.

**Input:** transcript-region raw current signal $s \in \mathbb{R}^T$ of one read; peak-distance $d_p$; slope-smoothing box width $w$; trim fraction $\tau$; pA outlier threshold $M$.
**Output:** ordered list of sub-events $\{(k, s_k, e_k, \mu_k, \sigma_k)\}$, where $[s_k, e_k)$ are sample boundaries and $(\mu_k, \sigma_k)$ are the in-segment statistics.

```
[Robust per-read normalization];
m ← median(s);   MAD ← median(|s − m|);   σ_r ← max(1.4826 · MAD, 10^−6);
s̃ ← (s − m)/σ_r ;                                  // robust z-score on the full read
[Boundary detection];
g ← |Δs̃| ;                                                        // absolute slope
ḡ ← boxsmooth(g, w);
P ← find_peaks(ḡ, distance = d_p);
if |P| ≥ 1 and P_0 < d_p then
 | P ← P \ {P_0} ;                                      // drop leading short segment
end
B ← (0, P_0, ..., P_{|P|−1}, T) ;                           // sub-event boundary list
[Per-sub-event statistics with τ-trimming];
out ← [];
for k ← 0 to |B| − 2 do
 | a ← B_k;   b ← B_{k+1};
 | if b ≤ a then
 | | continue
 | end
 | u ← s̃[a : b];
 | u′ ← trimboth(u, τ) ;                                      // drop τ from each end
 | μ_k ← mean(u′);   σ_k ← std(u′);
 | if ¬isfinite(μ_k) or |μ_k| > M or σ_k = 0 then
 | | μ_k ← NaN;   σ_k ← NaN ;                           // flag pathological segment
 | end
 | append (k, a, b, μ_k, σ_k) to out;
end
return out;
```

**Supplementary Algorithm S2:** Box-constrained anchored Viterbi alignment.

**Input:** sub-event means $\mathbf{e}_{1:N}$; per-position $k$-mer model $(\boldsymbol{\mu}_{1:L},\ \boldsymbol{\sigma}_{1:L})$; per-read shift $\beta$; emission offset $\varepsilon$ (precomputed once from $\boldsymbol{\sigma}$; see Algorithm S4); skip penalty $\lambda_{\text{skip}}$; boundary cost $\lambda_b$; per-sub-event length weights $\mathbf{w}_{1:N}$; basecalling anchors $(k_{\text{seed}},\ k_{\text{end}},\ j_{\text{min}},\ j_{\text{max}})$; entry/exit boxes $(e^{\ell}_{\text{st}}, e^{h}_{\text{st}}, e^{\ell}_{\text{en}}, e^{h}_{\text{en}}, k^{\ell}_{\text{st}}, k^{h}_{\text{st}}, k^{\ell}_{\text{en}}, k^{h}_{\text{en}})$; $k$-mer band $(k^{\ell}_{\text{band}}, k^{h}_{\text{band}})$.

**Output:** optimal start/end coordinates $(i^*,\ j^*)$, alignment log-likelihood $\ell^*$, and per-sub-event alignment vector $\mathbf{a} \in \mathbb{Z}^N$ (with $a_u = -1$ for unaligned).

set $j_{\text{exit}} \leftarrow j_{\max} + 1$; initialize $V \leftarrow -\infty$ and $B \leftarrow 0$;
`//` $B$`: 0=match, 1=stay, 2=skip, 3=entry-init sentinel;`
[**Entry-box initialization**];
**foreach** $(i, j)$ *in entry box* **do**
  $V[i,j] \leftarrow \max\big(V[i,j],\ -\lambda_b(\,|i - k_{\text{seed}}| + |j - j_{\min}|\,)\big)$;  $B[i,j] \leftarrow 3$;
**end**
[**Forward DP within $k$-mer band**];
**for** $i \leftarrow 1$ **to** $N$ **do**
  $\tilde{e}_i \leftarrow e_i + \beta$;  $w_i \leftarrow \mathbf{w}[i-1]$;
  **for** $j \leftarrow \max(1, k^{\ell}_{\text{band}})$ **to** $\min(L, k^{h}_{\text{band}})$ **do**
    $\ell_{ij} \leftarrow w_i \cdot \big[\log \mathcal{N}(\tilde{e}_i \mid \mu_j,\ \sigma_j) + \varepsilon\big]$;
    $m \leftarrow \max\big(V[i{-}1, j{-}1] + \ell_{ij},\ \ V[i{-}1, j] + \ell_{ij},\ \ V[i, j{-}1] - \lambda_{\text{skip}}\big)$;
    **if** $m > V[i,j]$ **then**
      $V[i,j] \leftarrow m$;  update $B[i,j]$ to the argmax above ; `// commit only on improvement: preserves entry-box init (`$B$`=3); unreachable cells stay` $-\infty$
    **end**
  **end**
**end**
[**Exit-box search**];
(`found`, $s^*$) $\leftarrow$ (`false`, $-\infty$);
**foreach** $(i, j)$ *in exit box* **do**
  **if** $V[i,j]$ *reachable* **then**
    $s \leftarrow V[i,j] - \lambda_b(\,|i - k_{\text{end}}| + |j - j_{\text{exit}}|\,)$;
    **if** $\neg$ *found* ***or*** $s > s^*$ **then**
      $(i^*,\ j^*,\ s^*,\ \ell^*) \leftarrow (i,\ j,\ s,\ V[i,j])$;  `found` $\leftarrow$ `true`;
    **end**
  **end**
**end**
**if** $\neg$ *found* **then**
  **return** $(0,\ 0,\ -\infty,\ \mathbf{a} \equiv -1)$
**end**
[**Backtrack**];
$(i, j) \leftarrow (i^*, j^*)$;
**while** $i > 0$ ***and*** $j > 0$ ***and*** $V[i,j]$ *reachable* ***and*** $B[i,j] \neq 3$ **do**
  **if** $B[i,j] = 0$ **then**
    $a_i \leftarrow j - 1$;  $i \leftarrow i - 1$;  $j \leftarrow j - 1$
  **else if** $B[i,j] = 1$ **then**
    $a_i \leftarrow j - 1$;  $i \leftarrow i - 1$
  **else** `//` $B[i,j] = 2$`, skip`
    $j \leftarrow j - 1$
  **end**
**end**
**return** $(i^*,\ j^*,\ \ell^*,\ \mathbf{a})$;

**Supplementary Algorithm S3:** Per-read iterative alignment with closed-form shift refinement.

**Input:** sub-event means $\mathbf{e}$; position-specific table $(\boldsymbol{\mu}, \boldsymbol{\sigma})$; emission offset $\varepsilon$; scoring penalties $(\lambda_{\text{skip}}, \lambda_b)$; per-sub-event length weights $\mathbf{w}$; anchors $(k_{\text{seed}}, k_{\text{end}}, j_{\min}, j_{\max})$; relaxation radii $(\delta_e, \delta_k)$; shift bounds $[\beta_{\min}, \beta_{\max}]$; max inner iterations $T$; LL tolerance $\tau_\ell$.

**Output:** per-read alignment $\mathbf{a}$, final log-likelihood $\ell$, fitted shift $\beta$.

```
construct entry/exit boxes and the k-mer band by relaxing the anchors by (δ_e, δ_k) in both
  directions;
β ← 0;  ℓ_prev ← −∞;
for t ← 1 to T do
    (i*, j*, ℓ, a) ← AnchoredViterbi(e, μ, σ, β, ε, λ_skip, λ_b, w, anchors, boxes, band) ;
     // Algorithm S2
    valid ← {u : a_u ≥ 0};
    if |valid| ≥ 30 and t < T then
        β̂ ← median{ μ_{a_u} − e_u : u ∈ valid } ;          // closed-form robust shift
        β ← clip(β̂, β_min, β_max);
    end
    if t > 1 and |ℓ − ℓ_prev| < τ_ℓ then
        break ;                                             // early stop on LL convergence
    end
    ℓ_prev ← ℓ;
end
return (a, ℓ, β);
```

**Supplementary Algorithm S4:** Outer iteration for the position-specific $k$-mer parameter table on the control sample.

**Input:** set of usable control reads $\mathcal{R}$; ONT $k$-mer prior $\boldsymbol{\mu}^{\text{ONT}}$; per-position $\boldsymbol{\sigma}^{\text{ONT}}$ (held fixed throughout); prior weight $\kappa$; number of outer iterations $Q$; scoring penalties $(\lambda_{\text{skip}}, \lambda_b)$; per-read length weights $\{\mathbf{w}_r\}$ from segmentation.

**Output:** refined position-specific mean vector $\boldsymbol{\mu}^{\text{con}}$ and final per-read alignments.

initialize $\boldsymbol{\mu}^{\text{con}} \leftarrow \boldsymbol{\mu}^{\text{ONT}}$;

$\varepsilon \leftarrow \dfrac{1}{L}\sum_{j=1}^{L} \log \sigma_j^{\text{ONT}} + \frac{1}{2}\log(2\pi) + \frac{1}{2}$ ; `// emission offset;` $\boldsymbol{\sigma}$ `fixed` $\Rightarrow$ `computed once`

**for** $q \leftarrow 1$ **to** $Q$ **do**

  `// (a) align all reads under current table;`

  **foreach** *read* $r \in \mathcal{R}$ **do**

    $(\mathbf{a}_r, \ell_r, \beta_r) \leftarrow$ PerReadAlign($\mathbf{e}_r$, $\boldsymbol{\mu}^{\text{con}}$, $\boldsymbol{\sigma}^{\text{ONT}}$, $\varepsilon$, $\lambda_{\text{skip}}$, $\lambda_b$, $\mathbf{w}_r$) ; `// Algorithm S3`

  **end**

  `// (b) re-estimate per-position mean with Bayesian shrinkage;`

  initialize $\mathbf{s} \leftarrow \mathbf{0}$,;

  $\mathbf{c} \leftarrow \mathbf{0}$;

  **foreach** *read* $r \in \mathcal{R}$ *with valid alignment* **do**

    **foreach** *aligned pair* $(u, j)$ *in* $r$ **do**

      $z \leftarrow e_{r,u} + \beta_r$;

      $s_j \leftarrow s_j + z$;  $c_j \leftarrow c_j + 1$;

    **end**

  **end**

  **for** $j \leftarrow 1$ **to** $L$ **do**

    $\mu_j^{\text{con}} \leftarrow \dfrac{s_j + \kappa\, \mu_j^{\text{ONT}}}{c_j + \kappa}$ ; `//` $\kappa$ `anchors low-coverage positions`

  **end**

**end**

re-align all reads once more under the converged $\boldsymbol{\mu}^{\text{con}}$;

**return** $\boldsymbol{\mu}^{\text{con}}$ and per-read alignments;

### 4 Supplementary Methods

#### 4.1 Parameter settings across datasets of segSHAPE

Unless noted otherwise, all results were produced with the default parameters listed below, given per pipeline step from base-calling to structure prediction.

- **Base-calling.** Dorado in reference-guided mode with `--emit-moves --reference`. For RNA002, we use Dorado v0.9.6, model `rna002_70bps_hac@v3`; while for RNA004, we use Dorado v1.4.0, model `rna004_130bps_sup@v5.3.0`.
- **Read filtering and transcript-region extraction.** samtools v1.21 flag filter `-F 2324` (primary, forward-mapped); minimum reference-coverage fraction 0.8 (relaxed to 0.5 on short or truncated transcripts); `MAPQ` $\geq 20$; duplicate-read and split-read removal; contig keep-list `--keep-contig TETRA` for the smPC_002pool_wt and smPC_004pool_wt datasets.
- **Signal segmentation.** Per-read robust $z$-normalization by median and MAD (scaled by 1.4826); `scipy.signal.find_peaks` on the smoothed absolute signal slope with a fixed minimum peak-to-peak distance of 10 samples; per-sub-event mean and standard deviation from a 10% symmetric trimmed estimator (`--trim 0.1`).
- **Position-specific $k$-mer table.** $k = 5$ (RNA002) or $k = 9$ (RNA004); ONT-prior blend weight $\kappa = 50$ effective sub-events; $Q = 3$ outer refinement rounds; per-position standard deviation held fixed at the ONT/f5c prior, i.e. only the mean is updated.
- **Anchored Viterbi alignment.** Anchor relaxation $\delta_e = 50$ sub-events and $\delta_k = 15$ $k$-mers; constant skip penalty $\lambda_{\text{skip}} = 50$; zero boundary cost $\lambda_b = 0$ (so the entry/exit boxes act as hard windows rather than soft anchor-distance penalties); length-weighted emission with a capped per-sub-event weight $w_i = \min(\text{dwell}_i, 30)$ (the sub-event sample count, capped to tame stalls); emission offset $\varepsilon$ set automatically to $\frac{1}{L}\sum_j \log \sigma_j^{\text{con}} + \frac{1}{2}\log(2\pi) + \frac{1}{2}$; up to $R = 3$ inner baseline-shift refinement rounds with log-likelihood tolerance $10^{-1}$; per-read shift $\beta$ clipped to a chemistry-dependent envelope ($[-1, 1]$ for RNA002, $[-0.3, 0.3]$ for RNA004).
- **Modification scoring (default: Isolation Forest).** scikit-learn v1.3.0 `IsolationForest` with $M = 100$ trees, `max_samples='auto'`, `contamination=0.005`, `random_state=42`. The interchangeable GMM detector uses `GaussianMixture` with `covariance_type='full'`, `reg_covar=1e-6`, `random_state=42`, component count $K \in \{1, 2\}$ selected per position by BIC, and tail quantile $q = 0.005$; both tails correspond to the same 0.5% expected control-sample outlier rate.
- **Reactivity normalization and structure prediction.** Per-position $z$-score; smoothing window `--smooth-window 0` (no smoothing, by default); RNAfold v2.7.0 with `-p -d2 --noLP -P rna_andronescu2007.par --shapeMethod=D`; the centroid structure is the final segSHAPE prediction.

The entire upstream signal-processing pipeline (anchored alignment with basecalling prior, position-specific $k$-mer modeling, and per-read baseline shift correction) was applied with identical default parameters across all datasets in this study (Supplementary Table S1). The two downstream reactivity-processing parameters, the Isolation Forest `contamination` rate and the reactivity smoothing window, were likewise held at their defaults (`contamination = 0.005`, `--smooth-window 0`) for the three benchmark RNAs in Table 1 and for the RNA002 comparison against sm-PORE-cupine in Figure 5. For the TPP riboswitch demonstration we re-calibrated these two parameters to `contamination = 0.05` and `--smooth-window 5` to accommodate the substantially lower read coverage. The two adjustments act in complementary directions: a larger `contamination` value relaxes the per-position outlier threshold so that fewer modification calls are lost when per-position read support is sparse, while a non-zero smoothing window pools reactivity across neighboring positions to compensate for the increased per-position variance at low coverage. Neither parameter affects the upstream reactivity-ranking quality (the per-position single-stranded versus double-stranded discrimination measured by ROC-AUC), but only the absolute reactivity scale and its spatial smoothing prior to RNAfold's SHAPE pseudo-energy conversion. For the RNA004 comparison against sm-PORE-cupine in Figure 5, we relaxed the Isolation-Forest outlier threshold to `contamination = 0.02` to accommodate the sparser per-position read support on this chemistry.

#### 4.2 Details of benchmarking baselines

We benchmarked segSHAPE against six baselines: three sequence-only predictors and three reactivity-aware DRS pipelines. Below we give the exact tool versions, commands and parameters used for each; runnable reproduction scripts for the three DRS pipelines are provided in the `baselines/` directory of the code repository. All structure predictions, including those of the baselines, were evaluated on the RNAfold *centroid* structure for consistency with segSHAPE.

**Sequence-only predictors.** These fold the reference sequence without any reactivity input.

- **RNAPKplex** and **RNAfold** (ViennaRNA v2.7.0) [6]: run on the reference sequence alone with default settings (`RNAPKplex < reference.fa` and `RNAfold < reference.fa`).
- **MXfold2** [7]: run through the authors' web server (https://ws.sato-lab.org/mxfold2) on the bare reference sequence. MXfold2 imposes a 1000-nt input limit, so it could not be applied to the 1552-nt *B. subtilis* 16S rRNA.

**Reactivity-aware DRS pipelines.** Each was reproduced from its published code and run on the same RNA002 reads as segSHAPE.

- **nanoSHAPE** [2]: reads were resquiggled with Tombo, and modifications were called with `tombo detect_modifications model_sample_compare` (treated vs. control, `--fishers-method-context 1`). At each position the per-read statistics were Benjamini–Hochberg FDR-corrected and the reactivity was taken as the fraction of reads with adjusted $p < 0.05$ (positions with $< 100$ reads dropped). The profile was SHAPE-normalized by the standard 2–8% rule (exclude values above max(1.5 IQR, 90$^{\text{th}}$ percentile), then divide by the mean of the top 10%) and folded with RNAfold v2.7.0 (`-p -d2 --noLP -P rna_andronescu2007.par --shapeMethod=D`).
- **SMS-seq** [4]: a hard-constraint method rather than a continuous-reactivity one. From the same Tombo `model_sample_compare` per-read statistics, a read was counted as modified at a position when its Tombo statistic was $< 0.2$; positions with $\geq 100$ reads and a modification rate $> 0.4$ were emitted as forced single-stranded (`x`) constraints. Structures were predicted with RNAfold v2.7.0 in constraint mode (`-p -d2 --noLP -C`) using the built-in Turner 2004 parameters (no `-P`), matching the ViennaRNA web-server configuration used in the original study.
- **PORE-cupine** [3]: per-read events were extracted with Nanopolish `eventalign`, and for each reference position a one-class svm (`svm` from R `e1071`, v3.6.3, `type="one-classification"`, RBF kernel, `nu`= 0.001, `gamma`= 0.0009) was trained on the control current features (dwell and mean) and applied to the treated reads; the modification rate is the fraction of treated reads flagged as outliers. The reactivity was folded with RNAfold v2.7.0 (`-p -d2 --noLP -P rna_andronescu2007.par --shapeMethod=D`).

**sm-PORE-cupine (RNA004 comparison).** For the RNA004 benchmark we compared against sm-PORE-cupine [5] rather than re-running it: the per-position modification rates on both RNA002 and RNA004 chemistries were downloaded directly from the accompanying Code Ocean capsule (https://codeocean.com/capsule/2426344/tree/v2), $z$-score normalized, and folded with the same RNAfold settings as segSHAPE. The independent SAFA footprinting reference annotation used to score per-position reactivity (single- vs. double-stranded) was obtained from the Source Data of Extended Data Figure 6i in [5].

#### 4.3 Implementation of the per-position modification detectors

The ablation in Supplementary Table S3 compares seven per-position detectors that all consume the *same* aligned events produced by the segSHAPE upstream pipeline (anchored alignment with basecalling prior, position-specific $k$-mer model, per-read baseline shift correction) and differ only in how a modification rate is computed from the control event set $\mathbf{x}_i$ and the treated event set $\mathbf{y}_i$ at each reference position $i$. All seven are implemented in the `segshape mod-calling` step (module `segshape.reactivity.calling`) under `scikit-learn v1.3.0`; their exact definitions are:

- **Isolation Forest** (segSHAPE default): scikit-learn `IsolationForest` (`n_estimators`= 100, `contamination`= 0.005, fixed `random_state`) fitted on $\mathbf{x}_i$; the rate is the fraction of $\mathbf{y}_i$ predicted as outliers.
- **Gaussian mixture model** (segSHAPE alternative): a `GaussianMixture` (`covariance_type='full'`) is fitted on $\mathbf{x}_i$ with the component count $K \in \{1, 2\}$ chosen by the lower BIC; the cut-off is the 0.5% quantile of the control log-likelihood and the rate is the fraction of $\mathbf{y}_i$ below it.
- **One-Class SVM**: scikit-learn `OneClassSVM` (RBF kernel, `gamma='scale'`, `nu`= 0.005) fitted on $\mathbf{x}_i$; the rate is the fraction of $\mathbf{y}_i$ flagged as outliers.
- **Absolute median shift** (|Δmedian|, dmed): $\left|\,\text{median}(\mathbf{y}_i) - \text{median}(\mathbf{x}_i)\,\right|$.
- **Kolmogorov–Smirnov**: the two-sample KS statistic between $\mathbf{x}_i$ and $\mathbf{y}_i$ (`scipy.stats.ks_2samp`).
- **Wasserstein**: the 1-D Wasserstein (earth-mover) distance between $\mathbf{x}_i$ and $\mathbf{y}_i$ (`scipy.stats.wasserstein_distance`).
- **xPore** [8]: a joint two-component Gaussian mixture is fitted to the pooled control and treated events by EM (component means initialized at the 25%/75% quantiles of the pooled data, the lower-mean component taken as unmodified); the rate is the treated posterior mass on the modified component.

The three threshold-based detectors (Isolation Forest, GMM, One-Class SVM) were calibrated to the same 0.5% expected control-sample outlier rate; the four distance/statistic detectors (|Δmedian|, KS, Wasserstein, xPore) are parameter-free.
